# Linking Individual Differences in Personalized Functional Network Topography to Psychopathology in Youth

**DOI:** 10.1101/2021.08.02.454763

**Authors:** Zaixu Cui, Adam R. Pines, Bart Larsen, Valerie J. Sydnor, Hongming Li, Azeez Adebimpe, Aaron F. Alexander-Bloch, Dani S. Bassett, Max Bertolero, Monica E. Calkins, Christos Davatzikos, Damien A. Fair, Ruben C. Gur, Raquel E. Gur, Tyler M. Moore, Sheila Shanmugan, Russell T. Shinohara, Jacob W. Vogel, Cedric H. Xia, Yong Fan, Theodore D. Satterthwaite

**Author notes:** Corresponding author. (Z.C.); (T.D.S.).

## Abstract

The spatial layout of large-scale functional brain networks differs between individuals and is particularly variable in association cortex that has been implicated in a broad range of psychiatric disorders. However, it remains unknown whether this variation in functional topography is related to major dimensions of psychopathology in youth. Capitalizing on a large sample with 27-minutes of high-quality functional MRI data (n=790, ages 8-23 years) and advances in machine learning, we examined associations between functional topography and four correlated dimensions of psychopathology (fear, psychosis, externalizing, anxious-misery) as well as an overall psychopathology factor. We found that functional topography significantly predicted individual differences in dimensions of psychopathology, driven mainly by robust associations between topography and overall psychopathology. Reduced cortical representations of association networks were among the most important features of the model. Our results emphasize the value of considering systematic differences in functional neuroanatomy for personalized diagnostics and therapeutics in psychiatry.

## INTRODUCTION

The human cerebral cortex is organized into spatially distributed large-scale functional networks that support diverse perceptual, executive, and socioemotional functions (Power et al., 2011; Yeo et al., 2011). Recent evidence from multiple independent studies has demonstrated that the spatial layout of these functional networks—their “functional topography”—varies substantially across individuals, even after accurate alignment to a common structural template (Bijsterbosch et al., 2018; Braga and Buckner, 2017; Glasser et al., 2016; Gordon et al., 2017a; Gordon et al., 2017b; Gordon et al., 2017c; Kong et al., 2019; Laumann et al., 2015; Li et al., 2019; Wang et al., 2015). Importantly, inter-individual variability in functional topography is not uniform across the cortex. Greater variability of functional topography is present in association cortex—higher-order, phylogenetically-expanded areas of cortex that support integrative, abstract, and advanced mental functions (Cui et al., 2020; Gordon et al., 2017b; Gordon et al., 2017c; Kong et al., 2019; Li et al., 2019; Wang et al., 2015). The functional topography of the association cortex is refined during development and has been previously associated with individual differences in executive function in youth (Cui et al., 2020). Critically, abnormal development of association cortices has been hypothesized to underlie the emergence of diverse psychiatric disorders during childhood, adolescence, and young adulthood (Menon, 2011; Sha et al., 2019; Sydnor et al., 2021). However, it remains unknown if individual differences in the functional topography of association cortex are linked to symptoms of psychiatric illness in youth.

While psychiatric illness is typically described according to the Diagnostic and Statistical Manual (DSM) (Edition, 2013), categorical clinical diagnoses fail to capture variability in disease severity, suffer from notable heterogeneity within diagnoses, and are marked by a high degree of co-morbidity (Clark et al., 2017; Drysdale et al., 2017; Kaczkurkin et al., 2020; Kotov et al., 2017; Lynch et al., 2020; Satterthwaite et al., 2020). Accordingly, efforts such as the National Institute of Mental Health (NIMH) Research Domain Criteria (RDoC) initiative and the Hierarchical Taxonomy of Psychopathology (HiTOP) frameworks have proposed dimensional models of psychopathology (Conway et al., 2019; Insel et al., 2010; Insel, 2014; Kotov et al., 2017; Krueger et al., 2018; Lahey et al., 2021). Dimensional taxonomies describe psychopathology as hierarchically organized, correlated dimensions of symptoms, wherein an individual receives a continuous score on each dimension (Kotov et al., 2017; Lahey et al., 2021). Previous studies conducted in youth samples have identified four major dimensions of psychopathology; these include fear, psychosis, externalizing and anxious-misery dimensions (Kaczkurkin et al., 2019; Kotov et al., 2017).

Emerging evidence additionally points to the importance of characterizing clinical and neural correlates of a dimensional *overall* psychopathology factor (also called the “*p*-factor”). The *p*-factor quantifies an individual’s shared vulnerability to a broad range of trans-diagnostic psychiatric symptoms and thus accounts for comorbidity among psychiatric disorders (Caspi et al., 2014; Caspi and Moffitt, 2018; Lahey et al., 2012; Lahey et al., 2017). Higher overall psychopathology scores, above and beyond specific psychopathology dimensions, are linked to earlier onset of psychiatric disorders and greater life impairment (Caspi et al., 2014). Moreover, using structural MRI, Romer and colleagues demonstrated that reduced cortical thickness commonly reported in association with specific psychiatric diagnoses may in fact represent a transdiagnostic feature associated with overall psychopathology severity (Romer et al., 2021). Notably, that study showed that reductions in thickness linked to psychopathology were largest in transmodal association cortices that sit at the top of a sensorimotor-association functional hierarchy (Romer et al., 2021). Similarly, using functional MRI, trans-diagnostic studies have reported dysfunction of association networks across diagnoses (Sha et al., 2019), which may reflect neurobiological correlates of pervasive comorbid psychopathology. Supporting this notion, the burden of overall psychopathology has been linked to abnormal patterns of functional connectivity between large-scale cortical networks (Elliott et al., 2018; Karcher et al., 2020; Kaufmann et al., 2017; Kebets et al., 2019). Nonetheless, it remains unknown whether individual differences in functional topography are related to major dimensions of psychopathology in youth, or whether abnormalities of association cortex topography are linked to overall psychopathology across disorders.

Here, we sought to understand how individual differences in functional network topography are associated with major dimensions of psychopathology in children, adolescents, and young adults. To do this, we capitalized on advances in machine learning and a large sample of youths who underwent clinical phenotyping and functional neuroimaging as part of the Philadelphia Neurodevelopmental Cohort (PNC) (Satterthwaite et al., 2014). We hypothesized that major dimensions of psychopathology would be linked to individual differences in functional topography. Furthermore, we expected that associations with major dimensions of psychopathology would be largely driven by shared deficits in association cortex linked to trans-diagnostic overall psychopathology.

## RESULTS

We studied 790 youths aged 8 to 23 years who underwent imaging as part of the PNC and had high-quality fMRI data of more than 27 minutes. As in our previous work (Cui et al., 2020), we used spatially regularized non-negative matrix factorization (NMF; Lee and Seung (1999)) to delineate 17 individual-specific (“personalized”) large-scale functional networks (Li et al., 2017) (**Figure S2**). NMF yields a probabilistic (soft) parcellation that can be converted into a discrete (hard) parcellation by labeling each vertex according to its highest loading (**Figure 1**). As previously (Cui et al., 2020), we named each personalized network according to its overlap with the canonical 17-network solution defined by Yeo *et al*. (Yeo et al., 2011). The spatial layout of these networks was largely similar to our prior work (Cui et al., 2020), with only subtle distinctions observed in the present sample, which includes persons with more servere psychopathology who were excluded from the previous study. As expected, across-subject variability of personalized functional network topography was highest in the association cortex (**Figure S3A and S3B**).

**Figure 1.**
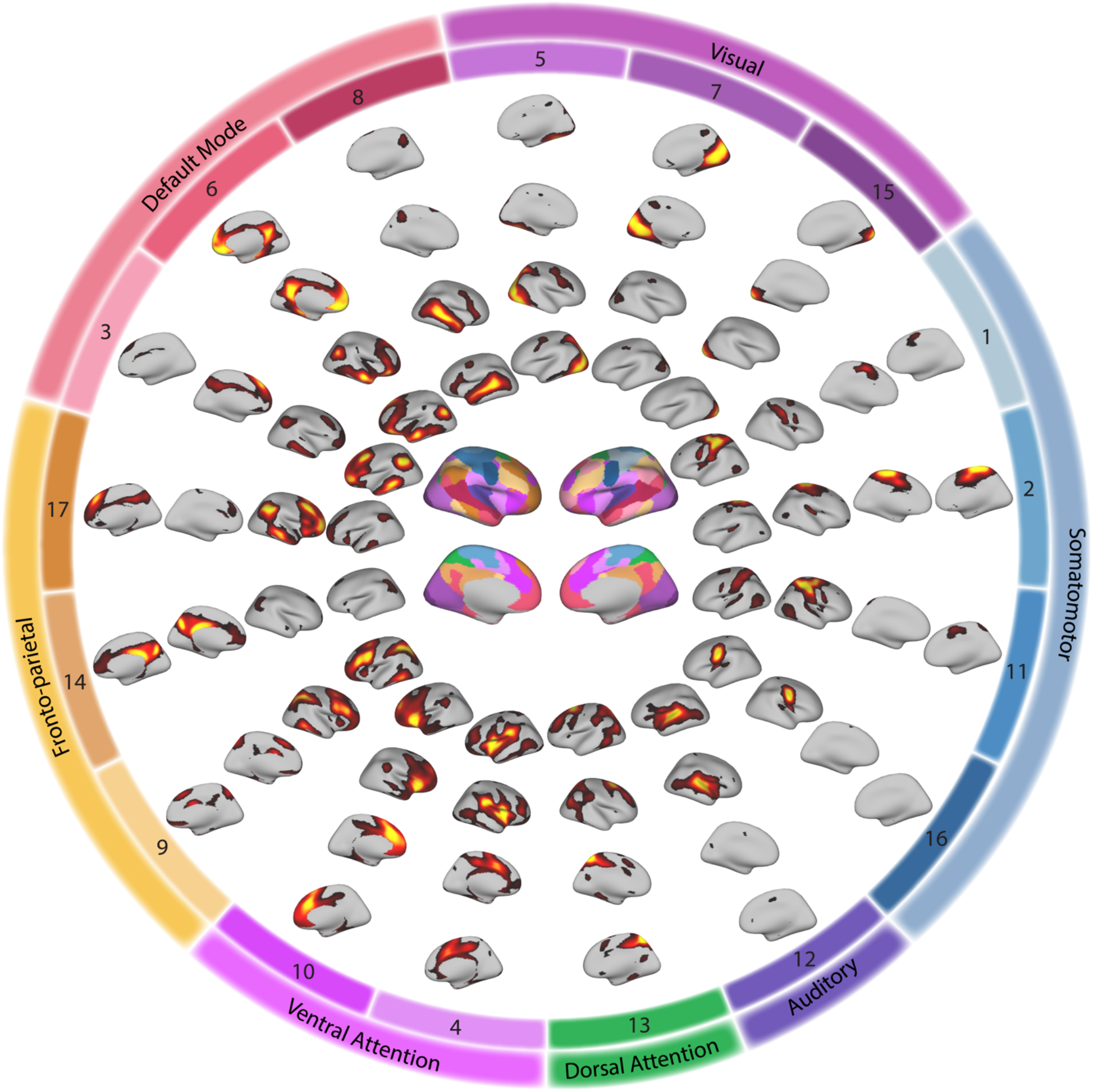
Group atlas used to initialize personalized functional networks. A group atlas was constructed to ensure correspondence across individuals; this group atlas was tailored to each individual to yield personalized networks. The networks in the group atlas include visual networks (5, 7, and 15), somatomotor networks (1, 2, 11, and 16), an auditory network (12), a dorsal attention network (13), ventral attention networks (4 and 10), fronto-parietal control networks (9, 14, and 17), and default mode networks (3, 6, and 8). In this atlas, there are 17 loadings for each vertex that quantify the extent to which the vertex belongs to each network. For each loading map, brighter colors indicate greater loadings. Vertices can be assigned to the network with the highest loading, yielding a discrete network parcellation (center).

### Functional topography predicts dimensions of psychopathology

We next related the topography of personalized networks to major dimensions of psychopathology. As described previously (Calkins et al., 2015; Calkins et al., 2017; Kaczkurkin et al., 2019; Shanmugan et al., 2016), psychopathology symptoms were assessed using a structured screening interview (GOASSESS) that included 112 items (see **Supplementary Data** and **Table S1)**. As in prior work (Kaczkurkin et al., 2019), we employed an exploratory factor analysis to create four correlated dimensions of psychopathology: fear, psychosis, externalizing, and anxious-misery (**Figure 2**). Each participant received a factor score for each of the four dimensions. The inter-factor correlations in this correlated-traits model were high (mean *r* = 0.71; see **Figure S4**). When dimensional scores were summarized by each psychiatric screening diagnosis, substantial similarities in the presence of each factor were observed across screening diagnoses (**Figure 2**).

**Figure 2.**
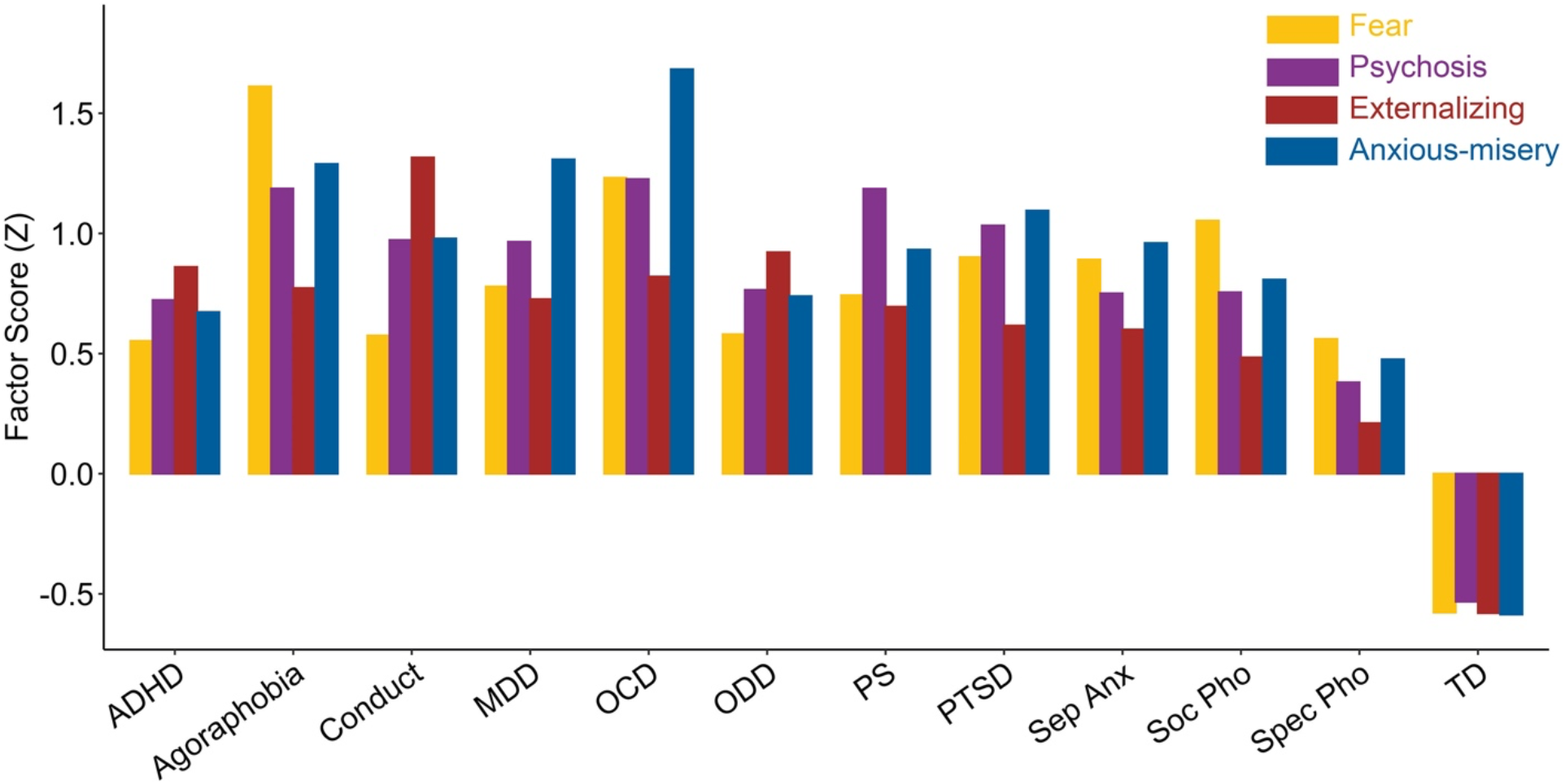
Correlated dimensions of psychopathology. An exploratory factor analysis of 112 psychopathology symptoms previously identified four correlated dimensions of psychopathology, including fear, psychosis, externalizing and anxious-misery. As expected, dimensional symptom profiles are substantially similar across screening diagnostic categories, as revealed by mean factor scores obtained for each category. See **Table S1** for the number of participants in each category. ADHD: attention deficit hyperactivity disorder; MDD: major depressive disorder; OCD: obsessive-compulsive disorder; ODD: oppositional defiant disorder; PS: psychosis spectrum; PTSD: post-traumatic stress disorder; Sep Anx: separation anxiety; Soc Pho: social phobia; Spec Pho: specific phobia; TD: typically developing. PNC: Philadelphia Neurodevelopmental Cohort.

Next, we investigated associations between factor scores and functional topography of personalized networks. Previous studies have demonstrated that multivariate analyses using machine learning can integrate spatially distributed predictive features in high-dimensional data (Haynes, 2015; Kong et al., 2019; Norman et al., 2006). Accordingly, we used a multivariate approach to understand the degree to which the overall pattern of functional topography encoded each of the four correlated dimensions of psychopathology. Specifically, we combined the loading maps of all 17 networks into a feature vector that represented each individual’s unique multivariate pattern of functional topography.

With these data, we used partial least square regression (PLS-R) to predict an individual’s score in each dimension of psychopathology from their functional topography. PLS-R seeks to find orthogonal latent components to predict an unseen outcome as accurately as possible; this approach is particularly suitable for high-dimensional data like functional topography. We used nested 2-fold cross-validation (2F-CV), with an outer loop evaluating the generalization of the model to unseen individuals, while the inner-loop selected optimal parameters (**Figure S5**). We evaluated each prediction by the correlation (*r*) between the actual and predicted scores, as well as by the mean absolute error (MAE). As the split into two halves was random, we repeated the above 2F-CV procedure 101 times and summarized the prediction accuracy using the median of the distribution. We used 101 repetitions rather than 100 to facilitate the selection of a median value. The significance of the prediction was evaluated using permutation testing, and Bonferroni correction was applied to account for multiple predictions (i.e., four dimensions). Throughout, we controlled for covariates including age, sex, and in-scanner motion.

This multivariate analysis revealed that the complex pattern of network topography could significantly predict unseen individuals’ dimensional scores of psychopathology. Specifically, we found that functional topography could predict symptoms of fear (*r* = 0.20, *P_Bonf_* < 0.001, MAE = 0.86, **Figure 3A**), psychosis (*r* = 0.16, *P_Bonf_* < 0.001, MAE = 0.89, **Figure 3B**), externalizing (*r* = 0.14, *P_Bonf_* < 0.001, MAE = 0.84, **Figure 3C**), and anxious-misery (*r* = 0.11, *P_Bonf_* = 0.008, MAE = 0.89 **Figure 3D**). These Bonferroni-corrected permutation tests indicate that the correlation between the actual and the PLS-R predicted scores was significantly higher than expected by chance for each of the four dimensions (**Figure 3E**).

**Figure 3.**
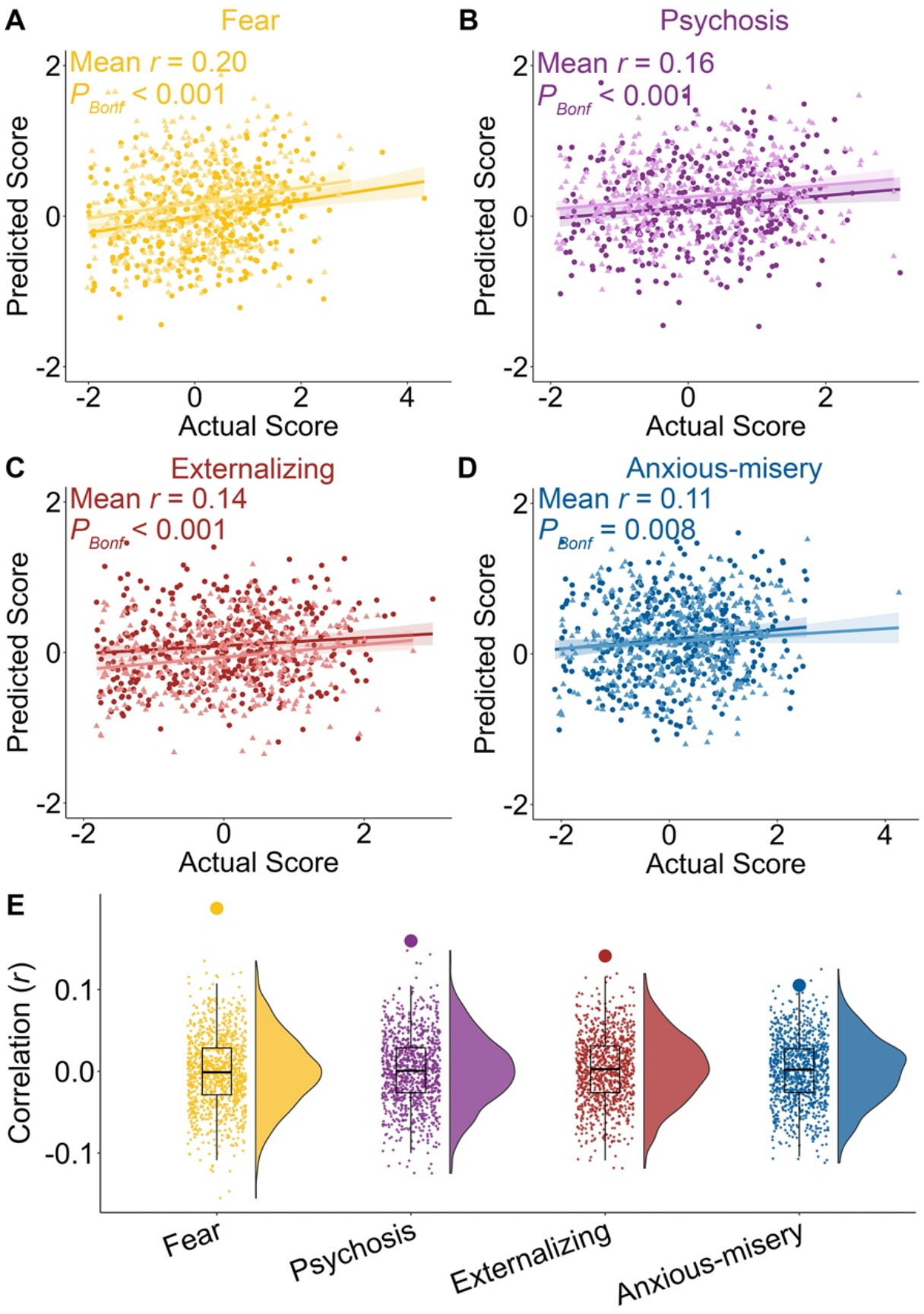
Functional topography predicts individual differences in major dimensions of psychopathology. Functional topography predicts unseen individuals’ dimensions of psychopathology, including fear (**A**), psychosis (**B**), externalizing (**C**) and anxious-misery (**D**). The data points represent the predicted scores (y-axis) of participants in a model trained on independent data using two-fold cross-validation (2F-CV). The 2F-CV was implemented by splitting all participants into two subsets. In each panel, the dark and light color represents participants of the two subsets, respectively. *P* values derived from permutation testing with Bonferroni correction indicated that the actual prediction accuracy (i.e., mean correlation *r* between two folds) was significantly higher than that expected by chance for all the four dimensions. Panel (**E**) shows the distribution of prediction accuracy values (i.e., correlation *r*) from permutation testing (small dots and histogram/boxplot) and the actual prediction accuracy (large dot).

### Major dimensions of psychopathology are predicted by similar patterns of functional topography

To understand the underlying patterns of network topography that contributed to our multivariate models, we evaluated the contribution weights of model features. In each multivariate model, every vertex received a feature weight for each network (i.e., 17 values per vertex). The absolute value of the weight quantifies the importance of the feature in the predictive model, while the sign indicates a negative or positive association between network loading and the dimensional psychopathology factor scores. Across the 101 split-half runs, we evaluated the median weight for each feature to summarize the contribution weight. Finally, to derive a network-level summary measure, we summed all feature weights within each network; this measure provides an interpretable summary of high-dimensional feature weights. A positive network-level summary value would indicate that a network has greater representation in individuals with more symptoms. In contrast, a negative network-level summary value would indicate that a network has a reduced representation in individuals with more symptoms.

This procedure revealed that deficits in association networks drove the prediction of each dimension of psychopathology, suggesting that there is reduced cortical representation of higher-order functional networks in youth with more severe psychiatric symptoms (**Figure 4**). For example, feature weights in fronto-parietal networks (networks 9 and 17) were negative in models of each of the four correlated dimensions of psychopathology (**Figure 4A-D**). Additionally, deficits in the ventral attention network (network 4) were prominent in psychosis (**Figure 4B**) and anxious-misery symptoms (**Figure 4D**), while deficits in the dorsal attention network (network 13) were notable for externalizing symptoms (**Figure 4C**). In contrast to findings in association cortex, positive feature weights were observed for lower-order somatomotor and visual networks, indicating a greater relative cortical representation of these networks in those with higher psychopathology symptoms. Together, these results suggest that deficits in the functional topography of association networks are linked to four correlated dimensions of psychopathology in youth.

**Figure 4.**
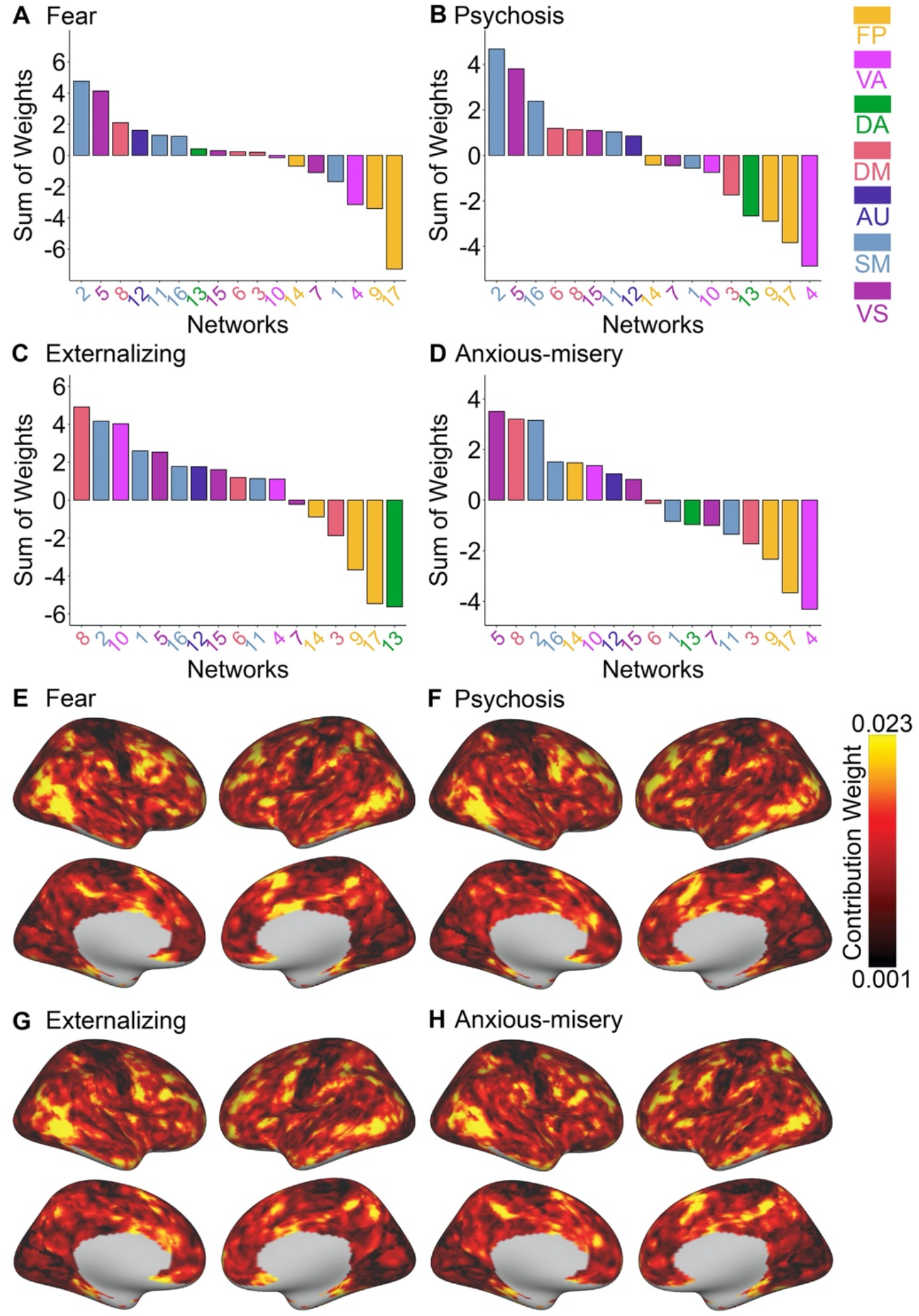
The four major dimensions of psychopathology are predicted by similar patterns of functional topography. Summing the model weights of all vertices within each network revealed that reduced cortical representation in association networks drove the prediction of fear (**A**), psychosis (**B**), externalizing (**C**) and anxious-misery (**D**) symptoms. At each location on the cortex, the absolute weight of each network was summed, revealing that the prefrontal, parietal, and occipital-temporal cortices contributed the most to the multivariate model in the prediction of fear (**E**), psychosis (**F**), externalizing (**G**) and anxious-misery (**H**) dimensions. FP: fronto-parietal; VA: ventral attention; DA: dorsal attention; DM: default mode; AU: auditory; SM: somatomotor; VS: visual.

While the above analysis summarized the contribution of features by network, we next sought to understand the location of important model features on the cortex. Accordingly, we examined the overall contribution of cortical locations in the multivariate model by summing the absolute weights of each vertex across all 17 networks. We observed that vertices in prefrontal, parietal, and temporo-occipital cortices contributed most to predicting the burden of psychopathology. This pattern was notably consistent across all four dimensions, including fear (**Figure 4E**), psychosis (**Figure 4F**), externalizing (**Figure 4G**), and anxious-misery (**Figure 4H**). To quantify the extent of similarity, we calculated the correlation of contribution patterns between each pair of dimensions, and assessed significance using a conservative spin-based spatial permutation procedure. This analysis revealed that the loading maps were similar across all dimensions (all pairwise *P_spin_ <* 0.001, mean pairwise *r* = 0.78; see **Figure S6**).

### Overall psychopathology underlies the similar functional topography pattern that predicts dimensions of psychopathology

The above analyses established that functional topography predicted the four correlated dimensions of psychopathology, and that the spatial distribution of predictive network patterns driving these effects were similar. These results prompted the hypothesis that these shared associations might be driven by a general psychopathology factor that captures symptoms integral to all four dimensions of psychopathology. To test this hypothesis, we used a previously-reported confirmatory bifactor analysis to parse the 112 item-level psychopathology symptoms into orthogonal dimensions of psychopathology (Kaczkurkin et al., 2019; Moore et al., 2019; Shanmugan et al., 2016). This model included five orthogonal dimensions of psychopathology, including overall psychopathology, fear, psychosis, externalizing, and anxious-misery (**Figure 5A**). Here, the overall psychopathology factor (also called the “*p*-factor”) describes a shared vulnerability to a broad range of symptoms across mental disorders (Caspi et al., 2014). Averaging factor scores by the screening diagnostic category revealed that overall psychopathology was high across all disorders. Further, after parsing out the effects of overall psychopathology, each disorder became more distinct in terms of the presence and severity of fear, psychosis, externalizing, and anxious-misery symptoms (**Figure 5B**).

**Figure 5.**
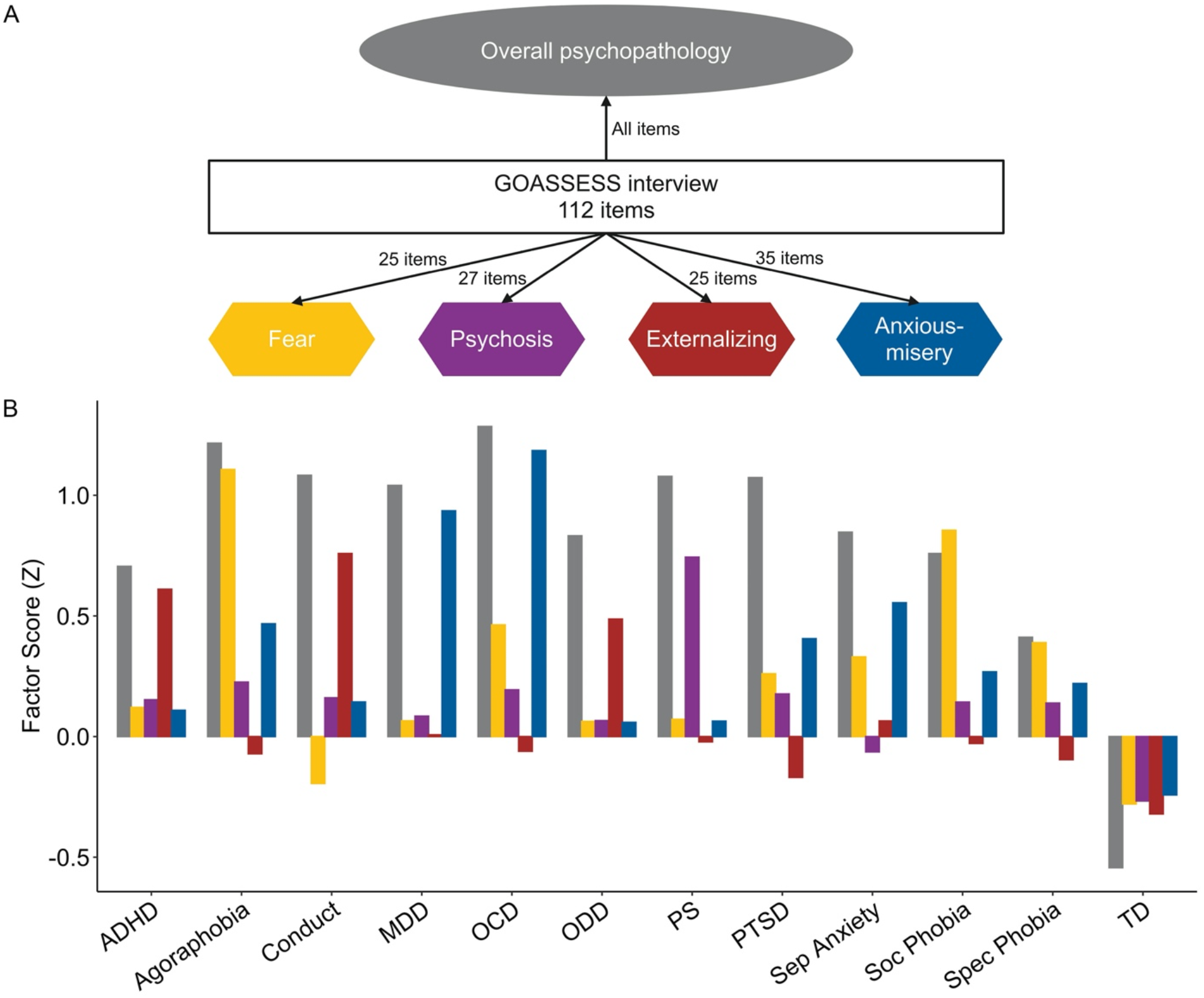
Common and divergent dimensions of psychopathology revealed by a bifactor model of psychopathology. (**A**) A confirmatory bifactor analysis was conducted on the 112 psychopathology items of the clinical screening interview to extract the orthogonal dimensions of psychopathology. These included four specific dimensions (i.e., fear, psychosis, externalizing and anxious-misery) and one common dimension (i.e., overall psychopathology). (**B**) The mean factor scores of the diagnostical categories illustrate that each specific psychopathology dimension loads more onto the relevant diagnostic categories, while the overall psychopathology factor loads onto all the diagnostic categories. ADHD: attention deficit hyperactivity disorder; MDD: major depressive disorder; OCD: obsessive-compulsive disorder; ODD: oppositional defiant disorder; PS: psychosis spectrum; PTSD: post-traumatic stress disorder; Sep Anx: separation anxiety; Soc Pho: Social Phobia; Spec Pho: Specific Phobia; TD: Typically developing.

To test whether the overall psychopathology factor drove the association between functional topography and correlated dimensions of psychopathology, we evaluated the degree to which the multivariate pattern of functional network topography could be used to predict unseen individuals’ scores of the overall psychopathology factor, using the same PLS-R prediction and permutation testing procedures as described above (e.g., **Figure S5**). We found that the multivariate pattern of functional topography significantly predicted overall psychopathology in unseen participants (*r* = 0.16, *P* < 0.001, MAE = 0.87; see **Figure 6A and 6B**). We next evaluated the features driving this model by network and by location on the cortex. We found that deficits (i.e. reduced cortical representation) of multiple association networks—including ventral attention (network 4), fronto-parietal (network 17), and dorsal attention (network 13) networks—drove the prediction of overall psychopathology (**Figure 6C**). In contrast, the overall contribution weights were positive in somatomotor (networks 2 and 16) and visual (network 5) networks (**Figure 6C**). To further examine these features, we evaluated the features with the highest (top 25%) absolute contribution weights (**Figure S7**). We observed that vertices in the ventral attention network (network 4; **Figure 6D**) and fronto-parietal network (e.g., network 17; **Figure 6E**) were predominantly assigned negative weights.

**Figure 6.**
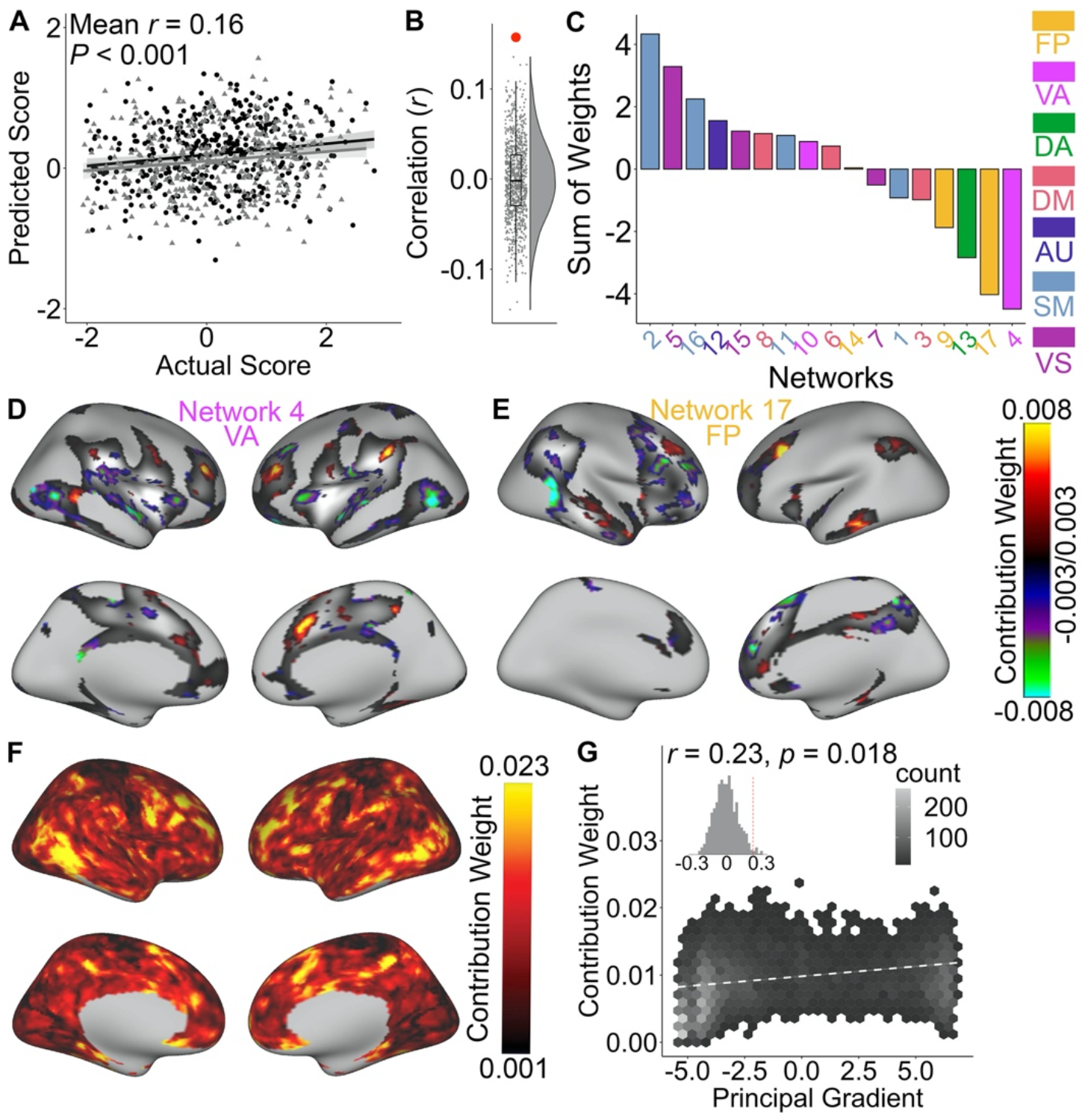
The functional topography of association networks predicts individual differences in the overall psychopathology factor. (**A**) Functional topography predicted unseen individuals’ overall psychopathology factor scores. Data points represent the predicted scores of the participants in a model trained on independent data using 2-fold cross-validation. The *P* value was derived from permutation testing. Panel (**B**) shows the distribution of prediction accuracies (i.e., correlation *r*) from permutation testing (small dots and histogram/boxplot) and the actual prediction accuracy (large red dot). (**C**) The fronto-parietal, ventral attention, and dorsal attention networks contained the highest negative contribution weights, indicative of an inverse relationship between the total cortical representation of those networks and overall psychopathology. (**D**) Model weights of features driving prediction mainly represented negative values in network 4, including the occipital-temporal junction, insula and inferior frontal areas. The top 25% of vertices in terms of feature importance are displayed. (**E**) The vertices in network 17 also mainly represented negative contribution weights in prefrontal areas and the occipital-temporal junction. (**F**) Vertices located at prefrontal, parietal, and occipital-temporal cortices drive the prediction of overall psychopathology. (**G**) The vertices that contributed the most were those sit at the top of principal gradient of functional connectivity (Margulies et al., 2016). FP: fronto-parietal; VA: ventral attention; DA: dorsal attention; DM: default mode; AU: auditory; SM: somatomotor; VS: visual.

Next, as previously, we summed the 17 absolute contribution weights for each vertex across networks, and found that the vertices in the prefrontal cortex and temporo-occipital junction contributed the most to the prediction of the overall psychopathology factor (**Figure 6F**). This contribution pattern aligned well (mean *r* = 0.86, all *P_spin_<* 0.001) with the patterns of contribution weights in the prediction models of the four correlated dimensions (see **Figure 4E-H**, above). This result suggests that, to a large extent, the association between functional topography and the overall psychopathology factor could explain the predictions of the four correlated dimensions of psychopathology. Finally, using spatial permutation testing, we evaluated the association between a vertex’s contribution to predicting overall psychopathology (**Figure 6F**) and its position along a sensorimotor-to-association cortical hierarchy defined by the principal gradient of functional connectivity in an independent dataset (**Figure S8**). This analysis revealed a significant positive correlation (*r* = 0.23, *P_spin_* = 0.018), indicating that transmodal regions in association cortex had higher feature weights in the model (**Figure 6G**).

Given the robust association identified between functional topography and overall psychopathology, we examined whether functional topography could predict other specific sub-factors of psychopathology from the bifactor model, which describe specific dimensions of psychopathology while accounting for overall psychopathology (**Figure 5**). Accordingly, we again used PLS-R and repeated (i.e., 101 times) nested 2F-CV as previously (**Figure S5**). We found that functional topography significantly predicted the fear factor (*r* = 0.11, *P_Bonf_* = 0.016, MAE = 0.89, **Figure S9A and S9B**). However, functional topography did not predict other specific dimensions from the bifactor model including psychosis (*r* = 0.08, *P_Bonf_* = 0.064, MAE = 0.87), anxious-misery (*r* = 0.08, *P_Bonf_* = 0.108, MAE = 0.87), or externalizing symptoms (*r* = 0.01, *P_Bonf_* = 1.000, MAE = 0.83). Notably, all prediction accuracies declined compared to those of the four correlated dimensions (**Figure 3**), suggesting that the association between functional topography and individual symptom dimensions was largely explained by overall psychopathology. Notably, we also observed both common and divergent patterns of functional topography explained the prediction of the fear factor and the overall psychopathology factor (**Supplementary Results**, **Figure S9**).

### Analysis of granular psychopathology symptoms provides convergent results

Overall, the above results demonstrated that the overall psychopathology factor dominated the association between functional topography and dimensional psychopathology. We next sought to further validate this result using an independent methodological approach. Specifically, instead of using the overall psychopathology index created by the bifactor model, we sought to find data-driven links between high-dimensional functional topography data and *item-level* symptoms of psychopathology from the structured screening interview (112 items). Accordingly, we used partial least square correlation (PLS-C), which finds pairs of latent components that maximize the correlation between two high-dimensional variables. As for our prior analyses, we used 2F-CVs to evaluate the generalizability of the correlation between pairs of latent components to unseen data. We repeated the 2F-CV procedure 101 times and summarized the out-of-sample correlation using the median of the distribution. Our analysis focused on the first pair of latent components, which explained the highest covariance (i.e., 5%) between topography and symptoms of psychopathology (**Figure S10**).

We found that the out-of-sample correlation between the first pair of latent components was significantly higher than that expected by chance (*r* = 0.18, *P*_perm_ < 0.001, **Figure 7A and 7B**), suggesting a pattern of association between a weighted set of psychopathology items and a weighted set of functional topographic features. As in previous studies (Griffis et al., 2019; Karlaftis et al., 2019), we examined the stability of the contribution weight of each psychopathology item to the first component. We found that *108* of the 112 psychopathology items contributed significantly to the first component (**Table S2**). Therefore, the first component represented a shared vulnerability to a broad range of symptoms, and thus reflected overall psychopathology.

**Figure 7.**
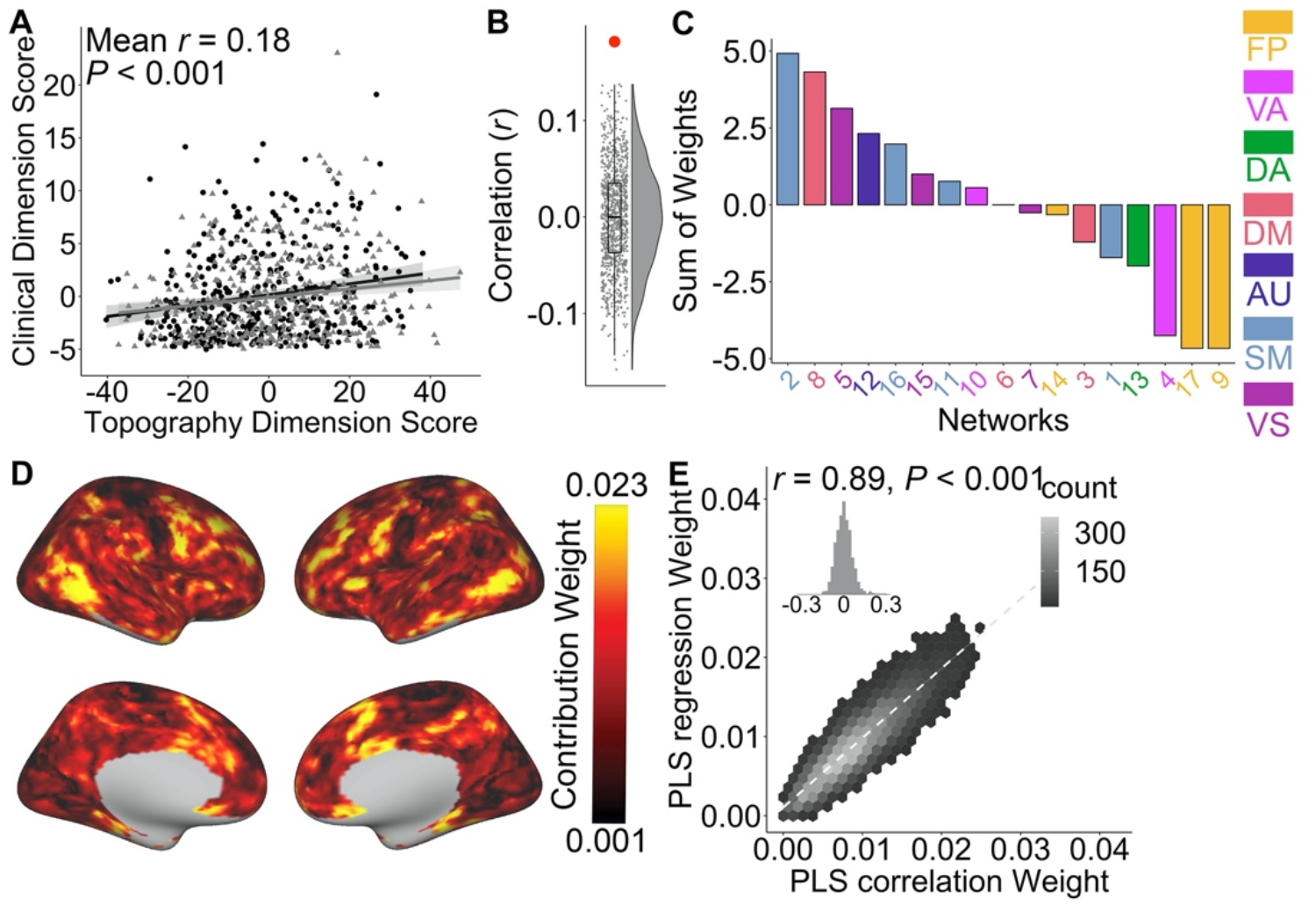
Linking the topography with *item-level* psychopathology symptoms revealed the association between functional topography and overall psychopathology factor. (**A**) The out-of-sample correlation between the first pair of topography and clinical dimension was *r* = 0.18 (*P*<0.001). Clinical dimension score was a weighted combination of all psychopathology items (See **Table S2** for weight of each item); while topography dimension score was a weighted combination of all network loadings. *P* value derived from permutation testing indicated that the actual out-of-sample correlation was significantly higher than that expected by chance. Panel (**B**) shows the distribution of out-of-sample correlation from permutation testing (small dots and histogram/boxplot) and the actual out-of-sample correlation (large red dot). (**C**) Examining the contribution weights of the topography pattern in the first latent component by summing the weights of all the vertices within each network revealed that the contribution weights were highly negative in the association networks, including fronto-parietal (networks 9 & 17) and ventral attention (network 4) networks. (**D**) At each location on the cortex, the absolute contribution weight of each network was summed, revealing that the prefrontal cortex and temporal-occipital junction contributed the most to the topography pattern in the first component. (**E**) This contribution pattern was highly correlated with the one (**Figure 6F**) from multivariate prediction model of overall psychopathology factor. FP: fronto-parietal; VA: ventral attention; DA: dorsal attention; DM: default mode; AU: auditory; SM: somatomotor; VS: visual.

Summing the contribution weights within each network, we found a convergent pattern of results from this data driven approach. Specifically, the networks with the most negative weights were in association cortex, including the fronto-parietal (networks 9 and 17) and ventral attention (network 4) networks (**Figure 7C**). In contrast, the somatomotor (network 2) and visual (network 5) networks had positive weights (**Figure 7C**). Summing the absolute contribution weight of each vertex across 17 networks indicated that the prefrontal cortex and temporal-occipital cortices contributed the most to this data-driven dimension that linked overall psychopathology and functional topography (**>Figure 7D**). Using spatial permutation testing, we found this cortical spatial distribution of contribution weights was strongly correlated (*r* = 0.89, *P_spin_* < 0.001, **Figure 7E**) with the contribution pattern from the prediction model of the overall psychopathology factor. In sum, this data-driven association between functional topography and *item-level* psychopathology symptoms provided convergent evidence that topography was related to overall psychopathology factor scores.

## DISCUSSION

We found that the spatial topography of large-scale, individual-specific functional networks was associated with individual differences in symptom severity within four major dimensions of psychopathology: fear, psychosis, externalizing and anxious-misery dimensions. Furthermore, we demonstrated that these associations between symptoms and functional topography were mainly driven by an individual’s level of overall psychopathology (commonly referred to as the *p*-factor). Critically, reduced cortical representation in association networks contributed the most to the prediction of overall psychopathology. Taken together, these findings suggest that individual differences in the spatial layout of association networks are systematically related to psychopathology in youth.

This work builds on convergent studies from multiple independent efforts that have demonstrated that there is marked individual variability in the topography of functional networks (Anderson et al., 2021; Bijsterbosch et al., 2018; Braga and Buckner, 2017; Cui et al., 2020; Glasser et al., 2016; Gordon et al., 2017a; Gordon et al., 2017b; Gordon et al., 2017c; Greene et al., 2020; Kong et al., 2019; Laumann et al., 2015; Li et al., 2019; Wang et al., 2015; Wang et al., 2018). Previous studies have reported that the individual variability of functional topography is maximum in association networks in adults (Cui et al., 2020; Gordon et al., 2017c; Kong et al., 2019; Wang et al., 2015); we previously found that as in adulthood, this is also true in childhood and adolescence (Cui et al., 2020). The present data further revealed the clinical relevance of this variability, by demonstrating that individual variation in functional topography is dimensionally associated with diverse forms of psychopathology in youth.

Current diagnostic systems (i.e., DSM) for psychiatric disorders assign patients into discrete categories based on signs and symptoms. However, it is increasingly recognized that these categories do not align with the underlying biology (Insel et al., 2010; Insel, 2014). Efforts including the RDoC initiative and HiTOP have been proposed given increasing empirical evidence that psychopathology is a dimensional phenomenon that is highly comorbid in nature, with the presence of one disorder increasing risk for the development of all others (Insel et al., 2010; Insel, 2014; Kotov et al., 2017; Kotov et al., 2021). In the current study, we used a dimensional approach that accounts for a continuous spectrum of psychopathology (Kaczkurkin et al., 2019). We identified four correlated major dimensions of psychopathology (i.e., fear, psychosis, externalizing and anxious-misery) that cut across the boundaries of traditional diagnoses. To parsimoniously account for shared vulnerability to all four symptom dimensions, we also used a bifactor model to identify four orthogonal dimensions and one overall psychopathology factor that reflects the shared burden of psychopathology across the four correlated dimensions (Caspi and Moffitt, 2018; Kaczkurkin et al., 2019; Lahey et al., 2021; Moore et al., 2019; Shanmugan et al., 2016).

We found that the topography of personalized functional networks significantly predicted the four major dimensions of psychopathology identified in our sample of youth. Although these dimensions are often considered independently, they are highly correlated. Perhaps as a result, we observed a substantial overlap between brain areas that strongly contributed to prediction across all four of the models, suggesting shared network contributions. We hypothesized that this overlap could be related to the clinical overlap among these dimensions, which reflects the high level of co-morbid psychopathology in individuals (Caspi and Moffitt, 2018; Lahey et al., 2021). Further analysis supported this interpretation by showing that the predictions were largely explained by the overall psychopathology factor. In particular, the contribution pattern predicting overall psychopathology overlapped with the features that predicted each of the four individual dimensions. Moreover, after accounting for the overall psychopathology factor, functional network topography no longer significantly predicted specific factors representing psychosis, externalizing, or anxious-misery dimensions. Together, these results suggest that individual variation in functional network topography may represent a broad vulnerability factor for trans-diagnostic psychopathology.

Consistent with our findings, previous transdiagnostic studies have reported common disrupted patterns of functional connectivity across mental disorders (Baker et al., 2019; Barron et al., 2020; Ma et al., 2020; Qi et al., 2020; Xia et al., 2018). Moreover, recent studies have demonstrated that the overall psychopathology factor is associated with abnormalities of both within- and between-network functional connectivity (Elliott et al., 2018; Karcher et al., 2020; Kebets et al., 2019). However, these studies calculated functional connectivity by applying a group-level functional atlas to each individual. Our results provide novel evidence that the spatial topography of the personalized functional networks is associated with the level of overall psychopathology. This is critical because topography and connectivity make distinct contributions to individual differences; using a group atlas for individuals’ data aliases the topographic signal into the measurement of connectivity (Bijsterbosch et al., 2018; Li et al., 2019). Future studies could leverage personalized functional network parcellations for the calculation of each individual’s functional connectivity to isolate the effects of both topography and connectivity (Gratton et al., 2019; Sylvester et al., 2020; Wang et al., 2018).

Importantly, we found that association networks, most notably ventral attention and fronto-parietal networks, predominantly displayed negative contribution weights in the multivariate prediction model of overall psychopathology severity, indicating that these networks were under-reprsented in youth with greater symptom burdens. Though our cross-sectional study cannot identify the underlying causal sequence, our results suggest that an individual with a reduced cortical representation of these executive networks is more likely to suffer from psychopathology. Consistent with our results, previous studies have demonstrated that dysfunction of the fronto-parietal (Baker et al., 2019; Cole et al., 2014; Xia et al., 2018) and ventral attention (Sheffield et al., 2017; Xia et al., 2018) networks is a common factor across a broad range of mental disorders. By summing the absolute contribution weights across the networks for each vertex, we observed that association cortex—which sits at the top of the sensorimotor-to-association cortical functional hierarchy (Margulies et al., 2016)—contributed the most to the prediction of overall psychopathology. This pattern may emerge given that association cortices support executive, social, and emotional mental functions implicated in psychopathology (Cole et al., 2014; Menon, 2011). Furthermore, the development of association networks is defined by a prolonged plastic period that enhances vulnerability to abnormal development and thus dysfunction (Larsen and Luna, 2018; Sydnor et al., 2021). Moreover, greater inter-individual variability observed in association networks nearer the top of the cortical hierarchy may contribute to the large variation in psychological functions subserved by association cortex in both healthy and psychiatric populations (Cui et al., 2020; Kong et al., 2019; Mueller et al., 2013; Wang et al., 2015).

Our prior study examining the refinement of functional network topography in youth established that the total cortical representation (relative extent) of individual association networks does not change over the developmental period of 8-23 years. Accordingly, the total amount of cortex that is occupied by each association network appears to be established earlier in life, during fetal, infant, or early childhood development. The relationship identified here between association network representation and overall psychopathology thus suggests that a shared vulnerability to diverse, comorbid, and severe psychiatric symptoms may be established early in life—with vulnerability transitioning to symptom manifestation throughout the course of the protracted development of association cortex. Critically, this may implicate developmental factors that are understood to determine the location and size of cortical functional areas in susceptibility to psychopathology, highlighting a path for future study. Such factors include patterning signaling molecules (e.g., fibroblast growth factors) that mechanistically regulate cortical arealization, as well as thalamo-cortical inputs that influence cortical area size and cortical properties during early development (Cadwell et al., 2019; Cholfin and Rubenstein, 2007; O’Leary et al., 2007).

In our study, several potential limitations and countervailing strengths should be noted. First, all data presented were cross-sectional, which precludes inferences regarding within-individual developmental effects. Studies with longitudinal sampling, such as the Adolescent Brain and Cognitive Development study (Casey et al., 2018) and the Human Connectome Project-Development (Somerville et al., 2018), will facilitate studies of within-individual changes of functional topography. Second, we combined data from the three fMRI runs, including two where an fMRI task was regressed from the data. This approach was motivated by prior evidence that functional networks are primarily defined by individual-specific rather than task-specific factors (Gratton et al., 2018), and that intrinsic networks during task performance are similar to those present at rest (Fair et al., 2007). Therefore, by including the task-regressed data, we were able to generate individualized networks using 27 min of high-quality data, which showed high concordance (*r* ∼ 0.92) with those generated using 380 min of data (Laumann et al., 2015). Third, certain important psychopathological classes, such as substance abuse, were not part of the screening interview and thus not included in the present analysis. Future studies may address this by considering broader assessments of psychopathology.

Despite these limitations, our study provides novel evidence that personalized functional network topography is related to overall psychopathology in youth. These findings emphasize the relevance of personalized functional neuroanatomy to the neurobiological mechanisms of comorbidity across psychiatric disorders. Because overall psychopathology in part explains a person’s liability to diverse symptoms of mental illness, the potential predictive power of functional network topography could potentially aid in the early identification of youths who are at risk of psychopathology. Finally, these results motivate clinical trials of neuromodulatory interventions that are targeted using personalized functional neuroanatomy.

## ACKNOWLEDGEMENTS

This study was supported by grants from the National Institute of Health: R01MH113550 (T.D.S. & D.S.B.), R01EB022573 (C.D., Y.F., & T.D.S.), R01MH120482 (T.D.S.), RF1MH116920 (T.D.S. & D.S.B.), R37MH125829 (D.F. & T.D.S), R01MH112847 (R.T.S. & T.D.S.), R01MH112070-01 (C.D.), R01MH096773 (D.A.F.), R01MH115357 (D.A.F.), R01MH119219 (R.C.G. & R.E.G.), R01NS060910 (R.T.S.), R01MH123563 (R.T.S.), F31MH123063-01 (A.P.), and T32MH014654 (B.L.). V.J.S. was supported by a National Science Foundation Graduate Research Fellowship (DGE-1845298). The PNC was supported by MH089983 and MH089924. Additional support was provided by the Lifespan Brain Institute at Penn and the Children’s Hospital of Philadelphia and the Dowshen Program for Neuroscience. The content is solely the responsibility of the authors and does not represent the official views of any of the funding agencies.

## AUTHOR CONTRIBUTIONS

T.D.S. and Z.C. designed the study. Z.C. and T.D.S. performed the analyses with support from A.R.P., H.L., and J.W.V.. H.L. and Y.F. provided parcellation tools. A.R.P. replicated all analyses. A.A., T.M.M., and T.D.S. completed data preprocessing. R.C.G., R.E.G., C.D., and T.D.S. provided resources. Z.C., A.R.P., B.L., V.J.S., and T.D.S. wrote the manuscript. H.L., A.A., A.F.A-B, D.S.B., M.B., M.E.C., C.D., D.A.F., R.C.G., R.E.G., T.M.M., S.S., R.T.S., J.W.V., C.H.X., and Y.F. reviewed and edited the manuscript.

## METHOD DETAILS

### Participants

Overall, 1,601 participants were studied as part of the PNC (Satterthwaite et al., 2014). However, 154 participants were excluded due to clinical factors, including medical disorders that could affect brain function or incidentally encountered structural brain abnormalities. Among the 1,447 subjects eligible for inclusion, 63 subjects were excluded for low quality in T1-weighted images or poor FreeSurfer reconstructions. Of the 1,384 subjects with usable T1 images and adequate FreeSurfer reconstructions, 594 participants were excluded for missing functional data or inadequate functional image quality; all participants were required to have three functional runs that passed quality assurance (QA). Specifically, as in prior work (Ciric et al., 2018; Satterthwaite et al., 2013a), a functional run was excluded if the mean relative root mean square (RMS) framewise displacement was higher than 0.2mm, or it had more than 20 frames with motion exceeding 0.25mm. This set of exclusion criteria resulted in a final sample of 790 participants (**Figure S1**), with a mean age of 16.04 years and a standard deviation (SD) of 3.21 years; the sample included 354 males and 436 females. All subjects or their parent/guardian provided informed consent, and minors provided assent. All study procedures were approved by the Institutional Review Boards of both the University of Pennsylvania and the Children’s Hospital of Philadelphia.

### Clinical assessment

Psychopathology symptoms were evaluated using a structured screening interview (GOASSESS), which has been described previously (Calkins et al., 2015; Calkins et al., 2017; Kaczkurkin et al., 2019; Shanmugan et al., 2016). GOASSESS is a structured screening interview based on a modified version of the *Kiddie-Schedule for Affective Disorders and Schizophrenia* (Kaufman et al., 1997) and *Diagnostic and Statistical Manual of Mental Disorders, 4^th^ edition,* Text Revision (Edition, 2013) criteria. In the PNC, the GOASSESS interview was administered by trained assessors who had undergone a common training protocol that included didactic sessions, assigned readings, and supervised observations. Before the experiment, all assessors were certified for conducting independent assessments, as evaluated by a 60-item checklist of interview procedures.

Before the experiment, all assessors had been certified for conducting independent assessments, as assessed using a 60-item checklist of interview procedures.

The GOASSESS psychopathology screen assesses the lifetime occurrence of psychosis spectrum symptoms, mood (major depressive episode and mania), anxiety (agoraphobia, generalized anxiety, panic, specific phobia, social phobia, separation anxiety, posttraumatic stress), behavioral disorders (oppositional defiant, attention deficit/hyperactivity, and conduct disorder), eating disorder (anorexia and bulimia), and suicidal thinking and behaviors. The 112 screening items (**Supplementary Data**) were administered to all participants. The frequency and relevant demographic data for each screening diagnosis considered are detailed in **Table S1**. Due to comorbidity, participants may be represented in more than one category. The median interval of time between clinical assessment and neuroimaging was 2 months.

### Exploratory Factor Analysis

As described elsewhere (Calkins et al., 2015; Kaczkurkin et al., 2018; Moore et al., 2019; Shanmugan et al., 2016), we conducted an exploratory factor analysis of the 112 item-level psychopathology symptoms based on the matrix of tetrachoric inter-item correlations (within the *psycho* package in R). We identified four correlated dimensions of psychopathology (**Figure 2**): fear (phobias), psychosis, externalizing, and anxious-misery (mood and anxiety).

### Confirmatory Factor analysis

Since the four dimensions acquired from the exploratory factor analysis were correlated, we hypothesized that there would be one general psychopathology factor (Caspi et al., 2014), which underlies the observed associations between these correlated dimensions. To test this hypothesis, we used a confirmatory bifactor analysis to parse the 112 item-level psychopathology symptoms into orthogonal dimensions of psychopathology (Calkins et al., 2015; Clark et al., 2021; Kaczkurkin et al., 2018; Moore et al., 2019; Shanmugan et al., 2016). The confirmatory bifactor model was estimated using a Bayesian estimator in Mplus (Muthen and Asparouhov, 2012). In a bifactor model, items load on up to two factors simultaneously, including their own “specific” factor and a general factor. The model generated five orthogonal dimensions of psychopathology (**Figure 5**): fear, psychosis, externalizing, anxious-misery, and overall psychopathology, the latter of which describes a shared vulnerability to a broad range of symptoms across mental disorders.

### Image acquisition

As previously described (Satterthwaite et al., 2014), all MRI scans were acquired using the same 3T Siemens Tim Trio whole-body scanner and 32-channel head coil at the Hospital of the University of Pennsylvania.

#### Structural MRI

Prior to the functional MRI acquisitions, a 5-min magnetization-prepared, rapid acquisition gradient-echo T1-weighted (MPRAGE) image (TR = 1810 ms; TE = 3.51 ms; TI = 1100 ms, FOV = 180 × 240 mm^2^, matrix = 192 × 256, effective voxel resolution = 0.9 × 0.9 × 1 mm^3^) was acquired.

#### Functional MRI

We used one resting-state and two task-based (i.e., *n*-back and emotion identification) fMRI scans as part of this study. All fMRI scans were acquired with the same single-shot, interleaved multi-slice, gradient-echo, echo planar imaging (GE-EPI) sequence sensitive to BOLD contrast with the following parameters: TR = 3000 ms; TE = 32 ms; flip angle = 90°; FOV = 192 × 192 mm^2^; matrix = 64 × 64; 46 slices; slice thickness/gap = 3/0 mm, effective voxel resolution = 3.0 × 3.0 × 3.0 mm^3^. Resting-state scans had 124 volumes, while the *n*-back and emotion recognition scans had 231 and 210 volumes, respectively. Further details regarding the *n*-back (Satterthwaite et al., 2013b) and emotion recognition (Wolf et al., 2015) tasks have been described in prior publications.

#### Field map

In addition, a B0 field map was derived for the application of distortion correction procedures, using a double-echo, gradient-recalled echo (GRE) sequence: TR = 1000ms; TE1 = 2.69ms; TE2 = 5.27ms; 44 slices; slice thickness/gap = 4/0 mm; FOV = 240 mm; effective voxel resolution = 3.8×3.8×4 mm.

#### Scanning procedure

To acclimate subjects to the MRI environment before scanning, a mock scanning session where subjects practiced the task was conducted using a decommissioned MRI scanner and head coil. Mock scanning was accompanied by acoustic recordings of the noise produced by gradient coils for each scanning pulse sequence. During these sessions, feedback regarding head movement was provided using the MoTrack motion tracking system (Psychology Software Tools). Motion feedback was given only during the mock scanning session. To further minimize participants’ motion, before data acquisition, participants’ head was stabilized in the head coil using one foam pad over each ear and a third over the top of the head.

### Image processing

The structural images were processed using FreeSurfer (version 5.3) to allow for the projection of functional time series to the cortical surface (Fischl, 2012). The functional images were processed using a top-performing preprocessing pipeline implemented via the eXtensible Connectivity Pipeline (XCP) Engine (Ciric et al., 2018), which includes tools from FSL (Jenkinson et al., 2012; Smith et al., 2004), AFNI (Cox, 1996) and ANTs (Avants et al., 2009). This pipeline included: (1) the correction for distortions induced by magnetic field inhomogeneity using FSL’s FUGUE utility; (2) the removal of the initial volumes of each acquisition (i.e., 4 volumes for the resting-state fMRI and 6 volumes for the emotion recognition task fMRI); (3) the realignment of all volumes to a selected reference volume using FSL’s MCFLIRT; (4) the interpolation of intensity outliers in each voxel’s time series using AFNI’s 3dDespike utility; (5) the demeaning and removal of any linear or quadratic trends; (6) the co-registration of functional data to the high-resolution structural image using boundary-based registration. The acquired images were de-noised through a 36-parameter confound regression model that has been shown to minimize associations with motion artifact while retaining the signals of interest in distinct sub-networks. This model included the six framewise estimates of motion, the mean signal extracted from eroded white matter and cerebrospinal fluid compartments, the mean signal extracted from the entire brain, as well as the derivatives of each of these nine aforementioned parameters, and quadratic terms of each of the nine parameters and their derivatives.

Both the BOLD-weighted time series and the artefactual model time series were temporally filtered using a first-order Butterworth filter with a passband between 0.01 and 0.08 HZ to avoid mismatch in the temporal domain (Hallquist et al., 2013). Furthermore, to derive “pseudo-resting state” time series that were comparable across runs, the task activation model was regressed from n-back or emotion identification fMRI data (Fair et al., 2007). The task activation model and nuisance matrix were regressed out using AFNI’s 3dTproject.

For each modality, the fMRI time series of each individual was projected to each subject’s FreeSurfer surface reconstruction and smoothed on the surface with a 6-mm full-width half-maximum (FWHM) kernel. The smoothed data was then projected to the *fsaverage5* template, which has 10,242 vertices on each hemisphere (18,715 vertices in total after removing the medial wall). Finally, we concatenated the three fMRI acquisitions, yielding 27 minutes and 45 seconds (555 time points) of functional data for each subject. As in the work from other groups (Gordon et al., 2016; Ojemann et al., 1997; Wig et al., 2014) and our prior work (Cui et al., 2020), we removed vertices with low signal-to-noise ratio (SNR), which mainly localized to the orbitofrontal cortex and ventral temporal cortex. The resulting inclusion mask consisted of 17,754 vertices.

### Regularized non-negative matrix factorization

As previously described (Li et al., 2017), we used non-negative matrix factorization (NMF; Lee and Seung (1999) to derive individualized functional networks. The NMF method factorizes the data by weighting cortical elements that positively covary, which leads to a highly specific and reproducible parts-based representation (Lee and Seung, 1999; Sotiras et al., 2017). Our approach was enhanced by a group consensus regularization term that preserves inter-individual correspondence, as well as a data locality regularization term that makes the decomposition robust to imaging noise, improves spatial smoothness, and enhances functional coherence of the subject-specific functional networks (see Li et al. (2017) for details of the method; see also: https://github.com/hmlicas/Collaborative_Brain_Decomposition). As NMF requires the input to be nonnegative values, we re-scaled the data by shifting time courses of each vertex linearly to ensure all values were positive (Li et al., 2017). To avoid features in greater numeric ranges dominating those in smaller numeric range, we further normalized the vertex-wise time course by its maximum value so that all the time points would have values in the range of [0, 1].

Given a group of *n* subjects, each having fMRI data *X^i^* ∈ *R*^×^, *i* = 1, …, *n*, consisting of *S* vertices and *T* time points, we aimed to find *K* non-negative functional networks 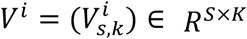 and their corresponding time courses 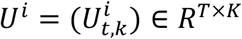 for each subject, such that

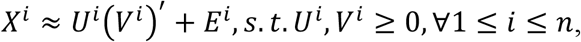

where (*V^i^*)′ is the transpose of (*V^i^*), and *E^i^* is independently and identically distributed (i.i.d) residual noise following Gaussian distribution with a probability density function of 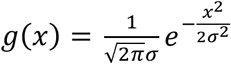. Both *U^i^* and *V^i^* were constrained to be non-negative so that each functional network does not contain any anti-correlated functional units (Lee and Seung, 1999). A group consensus regularization term was applied to ensure inter-individual correspondence, which was implemented as a scale-invariant group sparsity term on each column of *V^i^* and formulated as

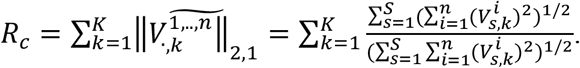

The data locality regularization term was applied to encourage spatial smoothness and coherence of the functional networks using graph regularization techniques (Cai et al., 2011). The data locality regularization term was formulated as

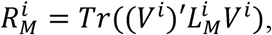

where 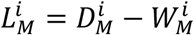 is a Laplacian matrix for subject *I*, 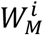 is a pairwise affinity matrix to measure spatial closeness or functional similarity between different vertices, and 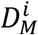 is its corresponding degree matrix. The affinity between each pair of spatially connected vertices (i.e., vertices *a* and *b*) was calculated as 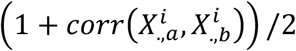, where 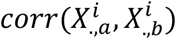 is the Pearson correlation coefficient between time series 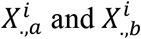, and others were set to zero so that 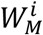 has a sparse structure. We identified subject-specific functional networks by optimizing a joint model with integrated data fitting and regularization terms formulated by

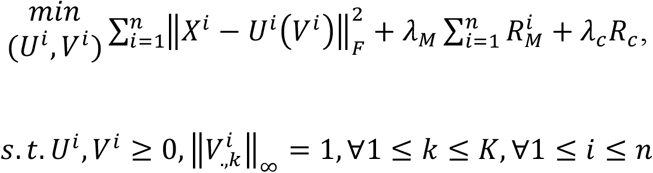

where 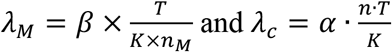 are used to balance the data fitting, data locality, and group consensus regularization terms, *n_M_* is the number of neighboring vertices, and *α* and *β* are free parameters. For this study, we used identical parameter settings as in prior validation studies (Li et al., 2017).

### Defining individualized networks

Our approach for defining individualized networks included three steps (see **Figure S2**). In the first two steps, a consensus group atlas was created. In the third step, this group atlas was used to define individualized networks for each participant. We decomposed the whole-brain into 17 networks, which allowed for a direct comparison to the other methods used in prior work (Kong et al., 2019; Wang et al., 2015; Yeo et al., 2011).

*Step 1: Group network initialization*. Although individuals exhibit distinct network topography, they are also broadly consistent (Gordon et al., 2017c; Gratton et al., 2018). Therefore, we first generated a group atlas and used it as an initialization for the definition of individualized networks. In this way, we also ensured spatial correspondence across all subjects. This strategy has also been applied in other methods for individualized network definitions (Kong et al., 2019; Wang et al., 2015). To avoid the group atlas being driven by outliers and to reduce the computational memory cost, a bootstrap strategy was utilized to perform the group-level decomposition multiple times on a subset of randomly selected subjects.

Subsequently, the resulting decomposition results were fused to obtain one robust initialization that is highly reproducible. As in our prior work (Cui et al., 2020; Li et al., 2017), we randomly selected 100 subjects and temporally concatenated their time series, resulting in a time series matrix with 55,500 rows (time-points) and 17,754 columns (vertices). Using a random non-negative initialization, we applied the above-mentioned regularized non-negative matrix factorization method to decompose this matrix into 17 functional networks (Lee and Seung, 1999). A group-level network loading matrix *V* was acquired, which had 17 rows and 17,754 columns. Each row of this matrix represented a functional network, while each column represented the loadings of a given cortical vertex. As in prior work (Li et al., 2017), this procedure was repeated 50 times, each time with a different subset of subjects; this procedure yielded 50 different group atlases.

*Step 2: Group network consensus*. Next, we combined the 50 group network atlases to obtain one robust and highly reproducible group network atlas using spectral clustering (Li et al., 2017). Specifically, we concatenated the 50 group parcellations together across networks and acquired a matrix with 850 rows (i.e., functional networks, abbreviated as FN) and 17,754 columns (i.e., vertices). The inter-network similarity was calculated as

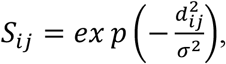

where *d_ij_* = 1 − *corr*(*FN_i_*, *FN_j_*), *corr*(*FN_i_*, *FN_j_*) is the Pearson correlation coefficient between *FN_i_* and *FN_j_*, and *σ* is the median of *d_ij_* across all possible pairs of FNs. Then, we applied the normalized-cuts (Cai et al., 2011) spectral clustering method to group the 850 FNs into 17 clusters. For each cluster, the FN with the highest overall similarity with all other FNs within the same cluster was selected as the most representative FN. The final group network atlas was composed of these maximally representative functional networks from each of the 17 clusters.

*Step 3: Individualized networks*. In this final step, we derived each individual’s specific network atlas using the acquired group networks (17 × 17,754 loading matrix) as the initialization and each individual’s specific fMRI times series (555 × 17,754 matrix). See (Li et al., 2017) for optimization details. This procedure yielded a loading matrix V (17 × 17,754 matrix) for each participant, where each row is a FN, each column is a vertex, and the value quantifies the extent to which each vertex belongs to each network. This probabilistic (soft) definition can be converted into discrete (hard) network definitions for display and comparison with other methods (Kong et al., 2019; Wang et al., 2015; Yeo et al., 2011) by labeling each vertex according to its highest loading.

### Across-subject variability of functional network topography

Prior studies have observed that across-subject variability of functional networks is higher in association networks and lower in primary sensorimotor networks in both adults (Gordon et al., 2017b; Gordon et al., 2017c; Kong et al., 2019; Li et al., 2019; Mueller et al., 2013; Wang et al., 2015) and youth (Cui et al., 2020). Here, we evaluated this observation in this sample, which included both healthy youths and youths with mental disorders. For each of the 17 networks, we calculated the median absolute deviation of loading values across all the subjects for each vertex. Next, we averaged the 17 median absolute deviation maps to generate the final across-network variability map that quantified the across-subject parcellation variance at each vertex.

### Prediction of psychopathology factors from functional network topography using partial least square regression (PLS-R)

We evaluated whether the multivariate spatial pattern of functional network topography encodes psychopathology. To address this question, we used partial least square regression (PLS-R) with nested two-fold cross validation (2F-CV, see **Figure S5**) to test whether multivariate network topography patterns could be used to identify the score of an unseen individual’s psychopathology factor in an unbiased fashion. We combined the loading maps of 17 networks into a *feature vector* to represent the multivariate spatial pattern of network topography of each individual. Then, we used these features to predict the four correlated dimensions of psychopathology (i.e., fear, psychosis, externalizing and anxious-misery) from the correlated traits model. Subsequently the same methods were used to predict each dimension (including overall psychopathology) from the bifactor model.

#### Partial least square regression (PLS-R)

PLS-R combines advantages from principal component analysis (PCA) and multiple linear regression to find orthogonal latent components to accurately predict outcomes. In contrast to other machine learning techniques (i.e., ridge regression) that using all features, PLS-R selected optimal latent components to do the prediction. Therefore, PLS-R could better suit in predicting dimensions of psychopathology, as prior studies have consistently demonstrated these dimensions are related to dissociated patterns of both brain structure and function (Kaczkurkin et al., 2019; Shanmugan et al., 2016; Xia et al., 2018). Here, we used the scikit-learn library to implement partial least square regression ((Pedregosa et al., 2011); algorithm ‘PLSRegression’ available at https://scikit-learn.org/stable/modules/cross_decomposition.html).

We set **X** to be a matrix of functional network topography with each row representing a participant and each column representing a *feature*, which was the network loading of one vertex in one network. We set **Y** to be a column vector of participants’ psychopathology factor scores. In contrast to PCA, which finds latent components to maximize the covariance within **X**, PLS-R searches for latent components that maximizes the covariance between **X** and **Y** by performing a double decomposition of **X** and **Y** using singular value decomposition (SVD). In this way, PLS-R seeks to find a set of orthogonal latent components of **X** that predict **Y** as accurate as possible (Abdi, 2003; Krishnan et al., 2011; Yoo et al., 2018).

PLS-R uses iterative applications of SVD to estimate latent components (Abdi, 2003). In each iteration, PLS-R finds the latent components of **X**, then subtracts the acquired latent components from the original **X** to yield the deflated matrix **X**_deflated_, which is used as the initial **X** for the next iteration. Similarly, the deflated matrix **Y**_deflated_ is updated by subtracting the corresponding predicted **Y** from the original **Y** in each iteration. We repeated this process until **X** was decomposed into *L* components. *L* was determined to be the optimal number of components via nested cross-validation (see the description in the *Prediction framework* below). During the optimization, PLS-R also fits the regression coefficients for each latent component to predict **Y** using multiple linear regression. By mapping these regression coefficients back to each functional topography feature (i.e., network loading), we generated one regression weight for each feature. The absolute values of these weights quantify the contribution of each functional topography feature to the prediction.

#### Prediction framework

We applied a nested 2-fold cross validation (2F-CV), with the outer 2F-CV estimating the generalizability of the model and the inner 2F-CV determining the optimal parameter (i.e., component number *L*) for the partial least square regression (See **Figure S5** for schematic of the prediction framework).

#### Outer 2F-CV

In the outer 2F-CV, the data was randomly divided into 2 subsets. We initially used subset 1 as the training set and subset 2 as the testing set. We regressed out age, sex and in-scanner head motion from each feature using a linear model on training data. To prevent leakage, the acquired beta coefficients were used to regress out these confounding factors from testing data. Then, each feature was linearly scaled between zero and one across the training dataset, and the acquired scaling parameters were also applied to scale the testing dataset (Cui and Gong, 2018; Cui et al., 2020). We applied an inner 2-fold cross validation (2F-CV) within the training set to select the optimal number (*L*) of components. Based on the optimal *L*, we trained a model using all subjects in the training set, and then used that model to predict the outcomes of all the subjects in the testing set. Analogously, we used subset 2 as the training set and subset 1 as the testing set, and repeated the above procedure. Across the testing subjects for each subset, the correlation and mean absolute error (MAE) between the predicted and actual outcomes were used to quantify the prediction accuracy. Both correlations and MAEs were averaged across two subsets to quantify the overall correlation and overall MAE.

#### Inner 2F-CV

Within each loop of the outer 2F-CV, we applied inner 2F-CVs to determine the optimal component number *L*. Specifically, the training set for each loop of the outer 2F-CV was further partitioned into 2 subsets randomly. One subset was selected to train the model under a given *L* in the range [1, 2, 3, …, 8, 9, 10], and the remaining subset was used to test the model. This procedure was repeated 2 times such that each subset would be used once as the testing dataset, resulting in two inner 2F-CV loops in total. For each value of *L*, the correlation *r* between the actual and the predicted outcome as well as the mean absolute error (MAE) were calculated for each inner 2F-CV loop, and then averaged across the two inner loops. The sum of the mean correlation *r* and reciprocal of the mean MAE was defined as the inner prediction accuracy, and the *L* with the highest inner prediction accuracy was chosen as the optimal parameter. Of note, the mean correlation *r* and the reciprocal of mean MAE cannot be summed directly, because the scales of the raw values of these two measures are quite different. Therefore, we normalized (using a *z*-score) the mean correlation *r* and the reciprocal of the mean MAE across all values and then summed the resultant normalized values.

Because the split was random, we repeated outer 2F-CV 101 times and calculated the median prediction accuracy (i.e., correlation *r* or MAE) across all 101 resamples to determine the overall prediction accuracy. We used 101 resamples rather than 100 to facilitate the selection of a median value. We visualized the scatter plot of the correlation between the predicted and actual scores for the repetition with median prediction accuracy. For computational efficiency, we executed the inner 2F-CV 20 times (Cui et al., 2020).

#### Significance of prediction performance

To evaluate whether prediction performance (i.e., correlation) was significantly better than expected by chance, we performed a permutation test (Mourao-Miranda et al., 2005). Specifically, the prediction procedure was re-applied 1,000 times. In each run, we permuted psychopathology factors across the training samples without replacement. Significance was determined by ranking the true prediction accuracy versus this permuted distribution. The *p-value* of the correlation *r* was the proportion of permutations that exhibited a higher value than the actual value (i.e., median correlation *r*) for the real data.

#### Interpreting model feature weights

From 101 repetitions of random 2F-CV, we acquired 202 regression coefficient or weight maps for each network. We calculated the median weight for each feature (i.e., loading of a vertex in one network), the absolute value of which quantifies the contribution of this feature to the model (Cui et al., 2020; Mourao-Miranda et al., 2005). To understand which network contributed the most to predictions, we summed the contribution weights across all vertices in each network. Finally, we calculated the spatial contribution of all vertices, in which we summed the absolute weight across all 17 networks to summarize the prediction weight of each vertex. This sum represented the importance of a given vertex to the predictive model.

### Linking functional network topography and pattern of psychopathology items using partial least square correlation (PLS-C)

Having demonstrated that the functional topography predicted unseen individual’s psychopathology *factor scores*, we further validated the association between topography and psychopathology by relating functional topography to *item-level* psychopathology data. In contrast to the analysis above (where clinical information was reduced to factor scores), here **Y** is a matrix with 790 rows and 112 columns, in which each row represents a participant and each column represents a single psychopathology item. We employed partial least square correlation (PLS-C) to examine the relationship between the two matrices (i.e., **X** and **Y**). Specifically, we used the scikit-learn library to implement partial least square regression (See Pedregosa et al. (2011); algorithm ‘PLSCanonical’ available at https://scikit-learn.org/stable/modules/cross_decomposition.html)

PLS-C also uses iterative applications of SVD to decompose the cross-product between functional topography matrix (**X)** and the *item-level* psychopathology matrix (**Y)**. In contrast to PLS-R, which aims to find the latent components of **X** to predict **Y**, PLS-C aims to find pairs of latent components *l***_X_** and *l***_Y_** with maximal covariance (Krishnan et al., 2011). Another difference from PLS-R, which updates the deflated matrix **Y**_deflated_ using the predicted **Y** in each iteration, is that PLS-C updates the deflated matrix **Y**_deflated_ using the latent components of **Y** (i.e., *l***_Y_**). The PLS-C model generated 112 pairs of latent components in total, as there were 112 psychopathology items. Each latent component represented a distinct pattern that relates a weighted set of psychopathology items to a weighted set of functional topography features. We calculated the median covariance explained by each component across repeated (i.e., 101) runs of 2F-CV. In our study, we focused on the first pair of latent components, which captured the highest and the most stable covariance (**Figure S10**). As in PLS-R analysis, we used repeated 2F-CVs (i.e., 101 repetitions) to evaluate the out-of-sample correlation of the first pair of latent components; we used the median correlation value across the 101 repetitions to summarize the overall association. Permutation testing (i.e., 1,000 times) was applied to test whether this correlation was significantly better than expected by chance.

As in prior work (Griffis et al., 2019; Karlaftis et al., 2019), we first evaluated the stability of the contribution weight of each psychopathology item to the first component to understand which items contributed to the multivariate model. As we repeated the 2-fold cross-validation 101 times, a total of 202 patterns of contribution weight were created for the first component. This procedure resulted in a distribution of contribution weight per psychopathology item. Random sampling can potentially induce arbitrary axis rotation, changing the ordering of canonical variates. Additionally, random sampling can induce axis reflection, which causes a sign change for the weights (Xia et al., 2018). In our data, the first pair of latent components was robust to arbitrary axis rotation, due to much higher **X**-**Y** covariance than the other components (**Figure S10**). To solve the axis reflection issue, we flipped the sign of the weight in all the psychopathology items and in all the functional topography features if the sign was negative in more than one half of psychopathology items. We calculated the *z*-score (i.e., mean/SD) of each psychopathology item across the 202 contribution weights. Psychopathology items with |*z*| > 2.576 were deemed to be significant contributors to the first component (Griffis et al., 2019; Karlaftis et al., 2019). We next sought to understand the contribution of functional topography to the first latent component. As PLS-R analysis, we acquired 202 weight maps and calculated the median weight for each vertex. We summed the contribution weights across all vertices in each network to understand which network contributed the most to predictions. Finally, we summed the absolute weight of each vertex across all 17 networks to summarize the spatial contribution of all vertices.

### Spatial permutation testing (spin test)

In order to evaluate the significance of the alignment among cortical contribution patterns as well as between contribution patterns and cortical functional hierarchy, we used a spatial permutation procedure called the spin test (Alexander-Bloch et al., 2018; Gordon et al., 2016; Sotiras et al., 2017; Vandekar et al., 2015) (https://github.com/spin-test/spin-test). The spin test is a spatial permutation method based on angular permutations of spherical projections at the cortical surface. Critically, the spin test preserves the spatial covariance structure of the data and consequently is far more conservative than randomly shuffling locations, which destroys the spatial covariance structure of the data and produces an unrealistically weak null distribution. In contrast, the spin test generates a null distribution of randomly rotated brain maps that preserve spatial features of the original map.

### Visualization

Connectome Workbench (version: 1.3.2) (Marcus et al., 2013); available at https://www.humanconnectome.org/software/connectome-workbench) was used to visualize the brain surface.

### Data & code availability

The PNC data is publicly available in the Database of Genotypes and Phenotypes: accession number: phs000607.v3.p2; https://www.ncbi.nlm.nih.gov/projects/gap/cgi-bin/study.cgi?study_id=phs000607.v3.p2. All analysis code is available here: https://github.com/ZaixuCui/pncsinglefuncparcel_psychopathology, with detailed explanation in https://zaixucui.github.io/pncsinglefuncparcel_psychopathology.

---

**Figure S1.**
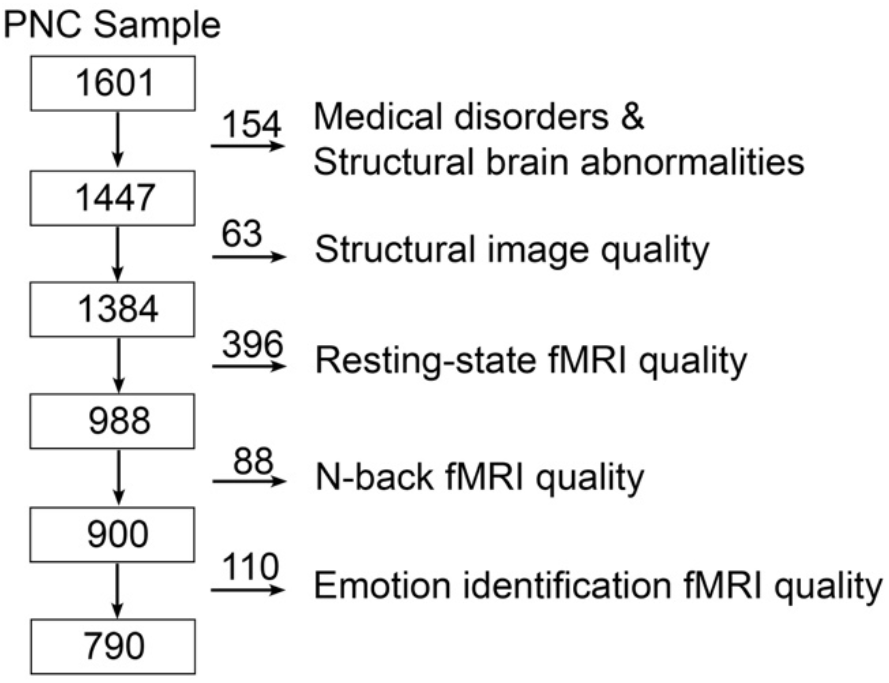
Sample construction. The cross-sectional sample of the Philadelphia Neurodevelopmental Cohort (PNC) includes 1601 participants in total. 154 participants were excluded because of clinical factors, including medical disorders that could affect brain function or incidentally encountered structural brain abnormalities. Then, 657 additional participants were excluded because of low quality of T1 or fMRI data. The final sample consisted of the remaining 790 participants.

**Figure S2.**
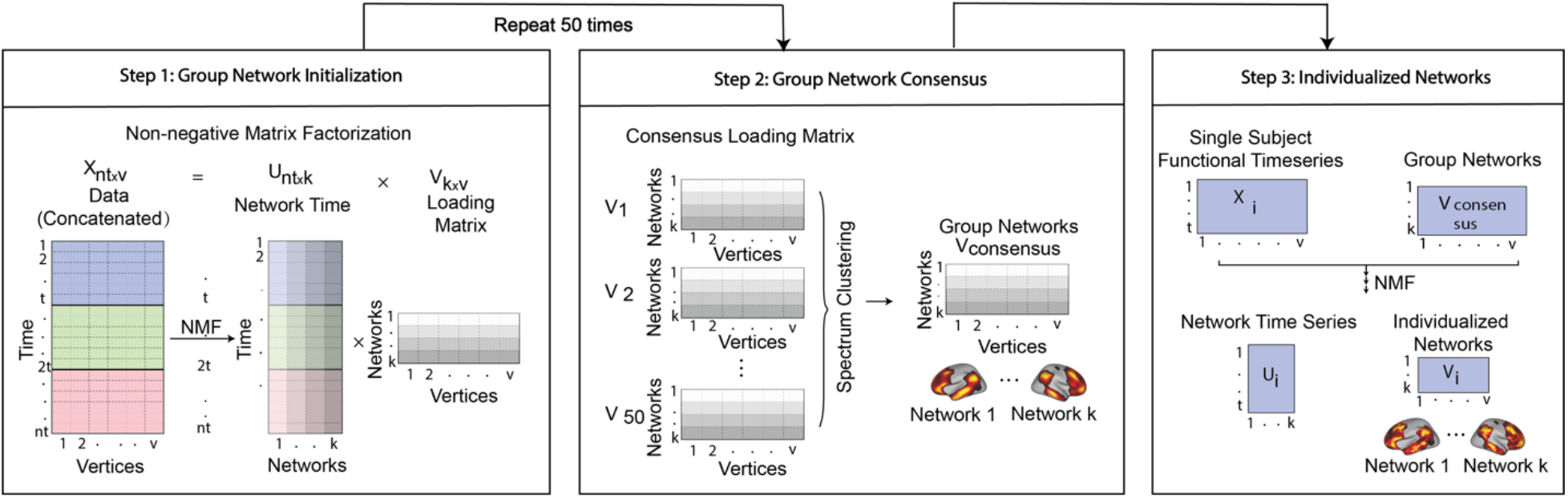
Schematic of spatially regularized non-negative matrix factorization (NMF) for individualized network parcellation. Related to STAR Methods. Each subject had three fMRI runs; we concatenated these for each subject, resulting in a 27.4-minutes time series with 555 time points for each subject. In the first step, we randomly selected 100 subjects and concatenated their time series into a matrix with 55,500 time points (rows) and 17,754 vertices (columns). Non-negative matrix factorization was used to decompose this data into a timeseries matrix and loading matrix. The loading matrix had 17 rows and 17,754 columns, which encoded the membership of each vertex for each network. This procedure was repeated 50 times, with each run including a different subset of 100 subjects. In the second step, a normalized-cut spectral clustering method was applied to cluster the 50 loading matrices into one consensus loading matrix, which served as the group atlas and ensured correspondence across individuals. In the third step, NMF was used to calculate individualized networks for each participant, with the group atlas used as a prior.

**Figure S3.**
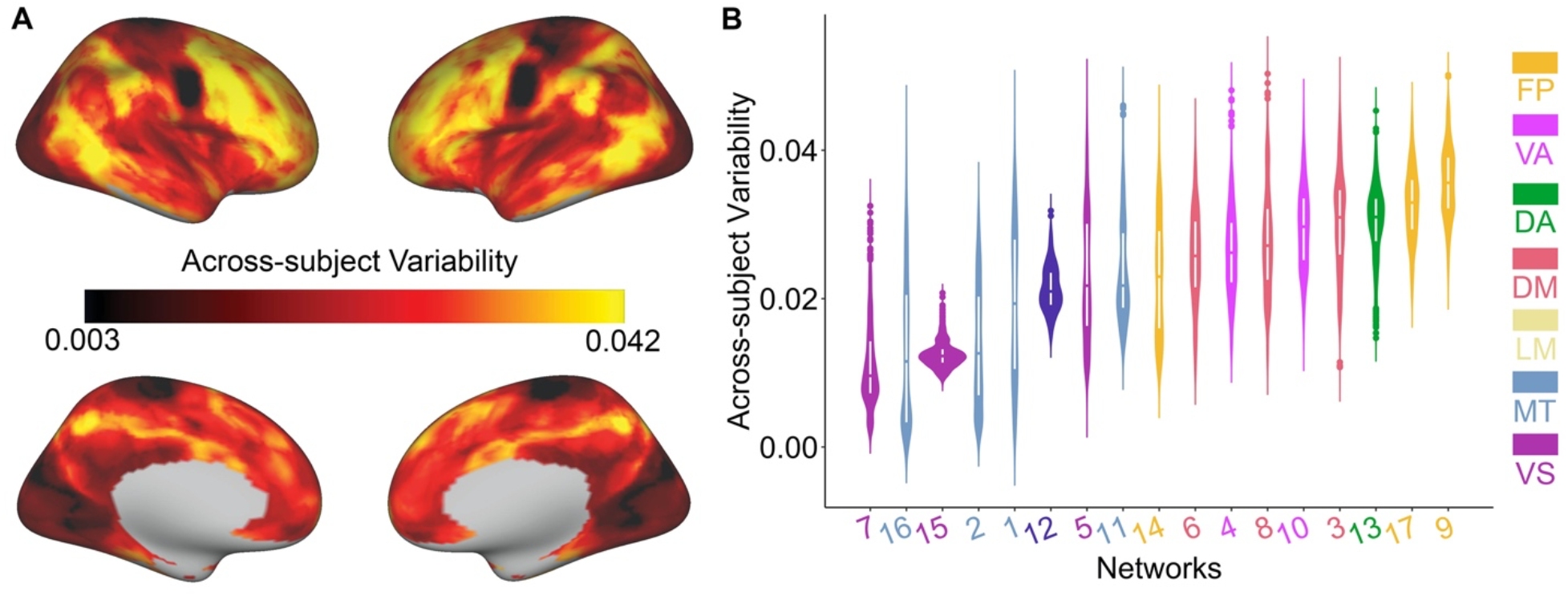
Across-subject variability of functional network topography is highest in association cortex. (**A**) A non-parametric measure of variability revealed that functional topography was most variable across individuals in association cortex and least variable in sensorimotor cortex. (**B**) Summarizing the cortical variability map by discrete functional networks confirmed that across-subject variability was highest in association networks, including the fronto-parietal, dorsal attention, default mode, and ventral attention networks.

**Figure S4.**
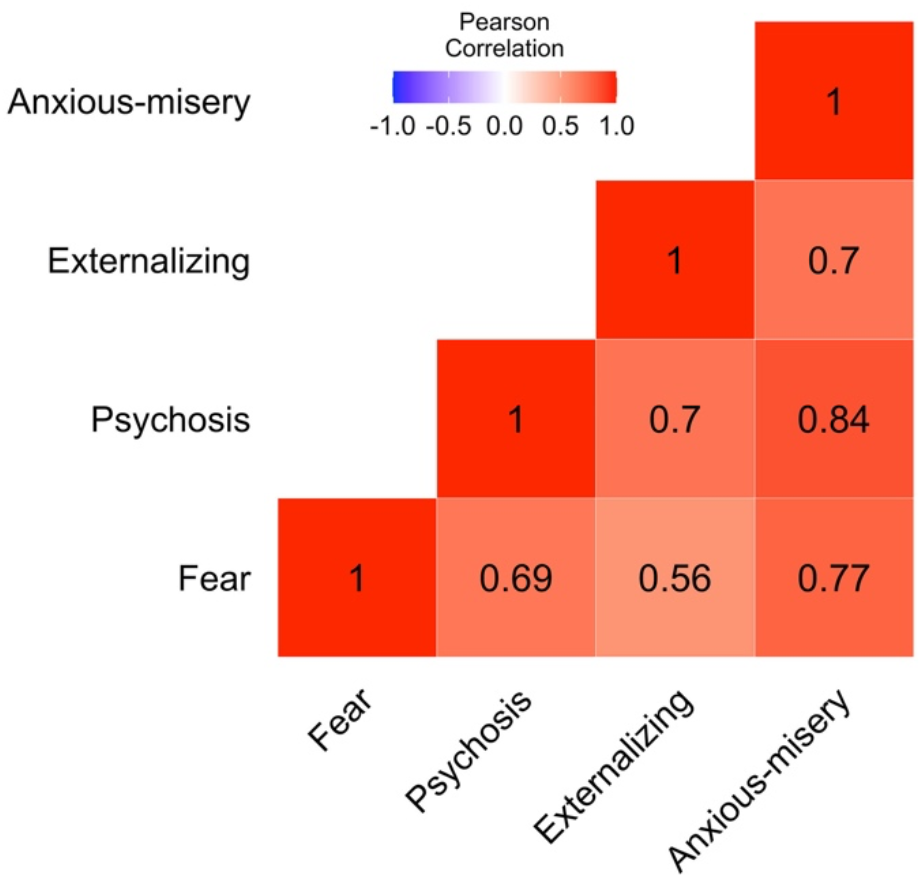
Factor scores of the four dimensions from the correlated traits exploratory factor model were correlated with each other.

**Figure S5.**
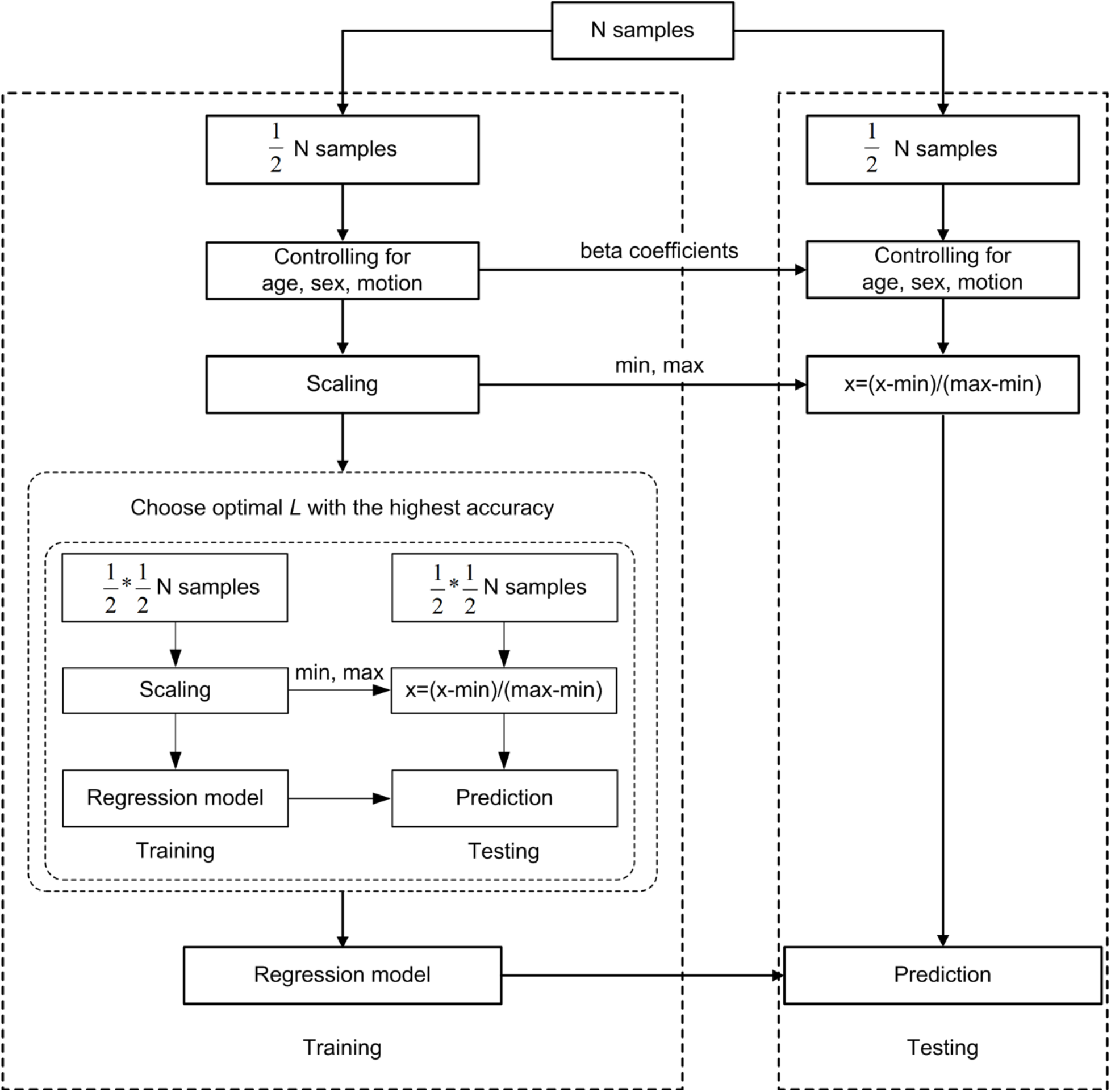
Schematic overview of one outer loop of the nested 2-fold cross-validation (2F-CV) prediction framework. All participants were divided into 2 halves randomly, with the first half used as a training set and the second half used as a testing set. We controlled for age, sex, and in-scanner motion from each feature in the training dataset, and then used the acquired coefficients to regress these covariates from the testing dataset. Each feature was linearly scaled between zero and one across the training dataset, and the scaling parameters were also applied to scale the testing dataset. An inner 2F-CV was applied within the training set to select the optimal parameter: component number *L*. Based on the optimal parameter, we trained a model using participants in the training set, and then used that model to predict the psychopathology scores of participants in the testing set. The prediction accuracy was evaluated by using the correlation r and mean absolute error between predicted and actual scores across participants in testing set.

**Figure S6.**
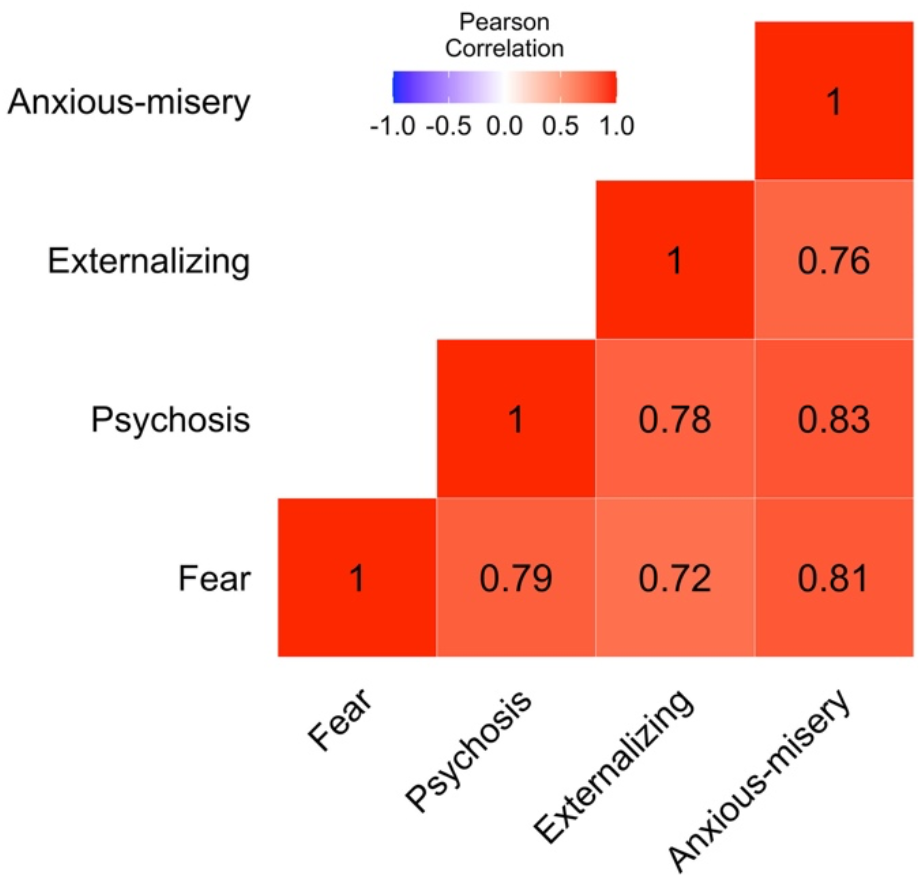
Feature weight maps from predictive analysis are highly correlated with each other. See Figure 4 E, F, G, H for the contribution maps of each of the four dimensions.

**Figure S7.**
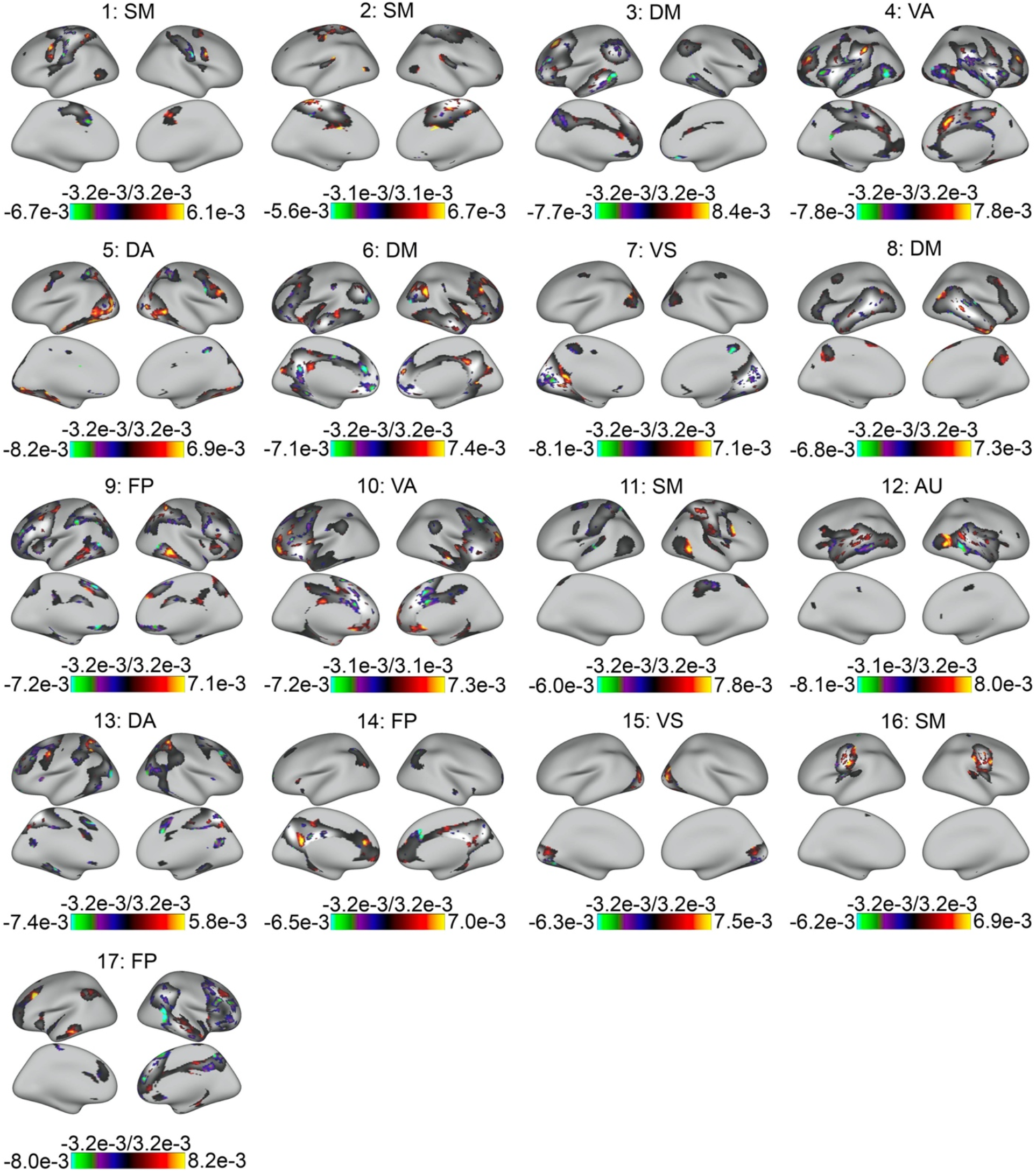
Features contributing to the multivariate prediction model of overall psychopathology. Contribution maps are thresholded to include vertices with the highest (first 25%) absolute contribution weight; underlay depicts the group network atlas. FP: fronto-parietal; VA: ventral attention; DA: dorsal attention; DM: default mode; AU: auditory; SM: somatomotor; VS: visual.

**Figure S8.**
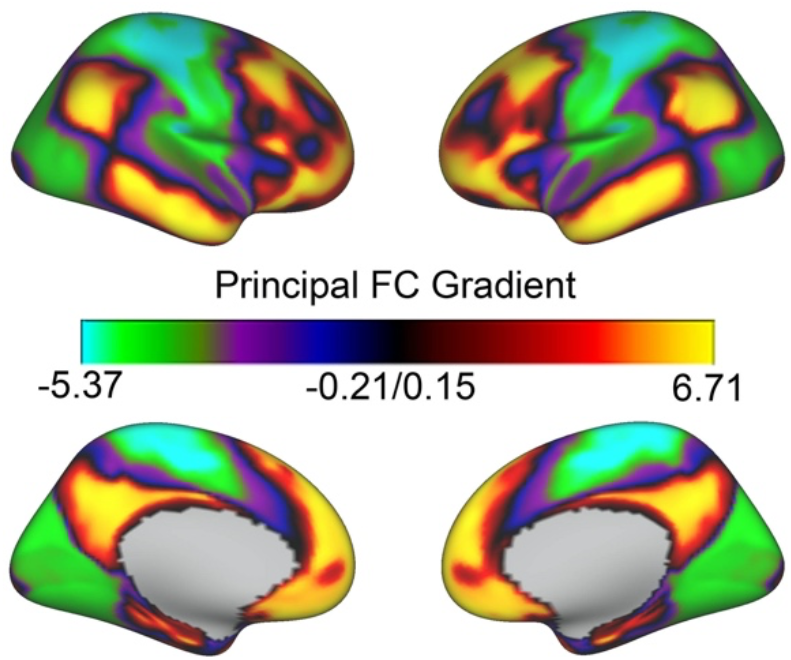
Principal gradient of functional connectivity from Margulies et al., 2016.

**Figure S9.**
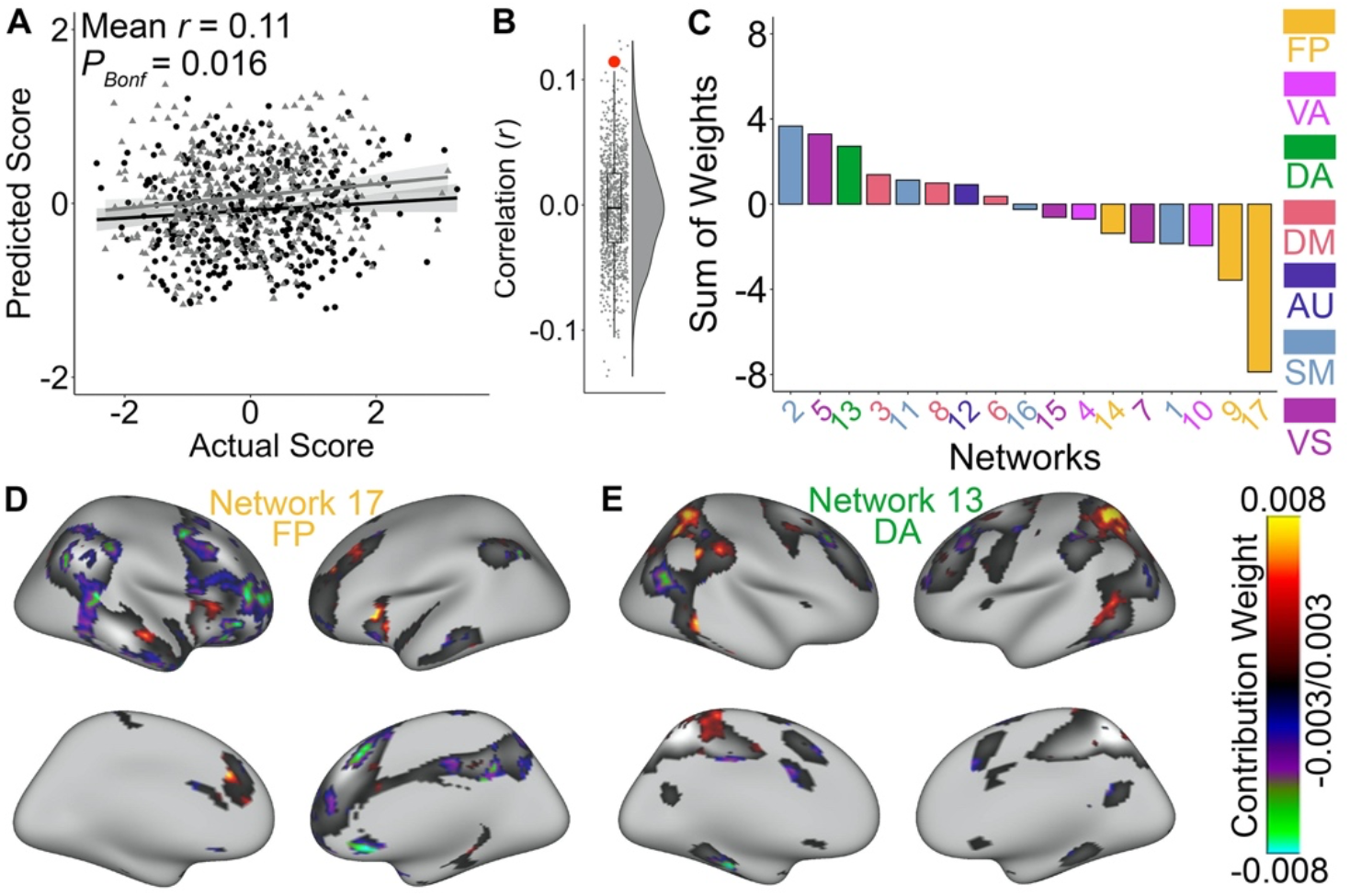
Patterns of functional topography that predict fear symptoms. (**A**) Functional topography predicted unseen individuals’ burden of specific fear symptoms quantified using the bifactor model. Data points represent the predicted scores of the participants in a model trained on independent data using 2-fold cross validation. *P* values derived from permutation testing with Bonferroni correction indicated that the prediction accuracy (i.e., correlation *r*) was significantly higher than that expected by chance. Panel (**B**) shows the distribution of prediction accuracies (i.e., correlation *r*) from permutation testing (small dots and histogram/boxplot) and the actual prediction accuracy (large red dot). (**C**) The fronto-parietal networks contained the greatest negative contribution weights, indicative of an inverse relationship between the total cortical representation of these networks and the fear factor. (**D**) The most highly weighted vertices in fronto-parietal network (i.e., network 17) mainly displayed negative contribution weights. (**E**) The most highly weighted vertices in dorsal attention network (i.e., network 13) mainly displayed positive contribution weights. FP: fronto-parietal; VA: ventral attention; DA: dorsal attention; DM: default mode; AU: auditory; SM: somatomotor; VS: visual.

**Figure S10.**
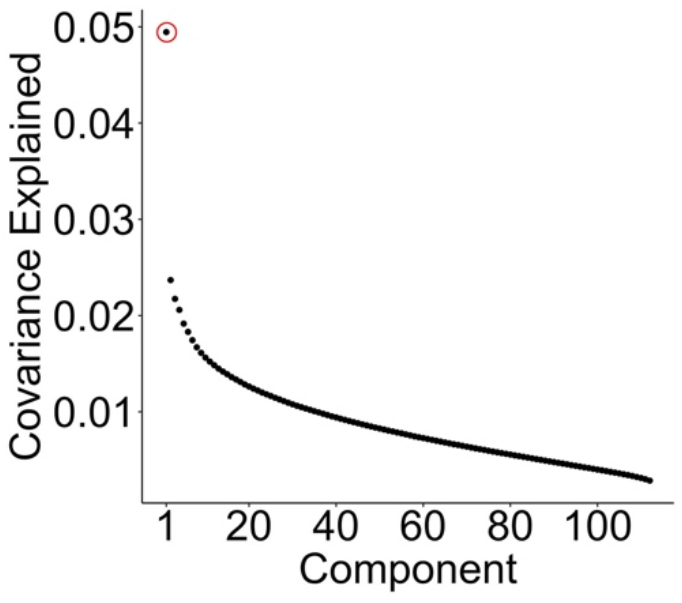
The first latent component explained the highest covariance (i.e., 5%) between functional topography and *item-level* psychopathology symptoms based on partial least square correlation analysis. The first component is labeled by a red circle.

**Table S1.** Summary of demographic and clinical data.

| Measure | Mean | SD |
| --- | --- | --- |
| Age (years) | 16.04 | 3.21 |
|  | N | % |
| Male | 354 | 45 |
| Lifetime prevalence<sup>a</sup> |  |  |
| Typically developing | 245 | 31 |
| Attention deficit hyperactivity disorder | 114 | 14 |
| Agoraphobia | 47 | 6 |
| Anorexia nervosa | 10 | 1 |
| Bulimia nervosa | 3 | <1 |
| Conduct disorder | 64 | 8 |
| Generalized anxiety disorder | 14 | 2 |
| Mania | 10 | 1 |
| Major depressive disorder | 132 | 17 |
| Obsessive-compulsive disorder | 21 | 3 |
| Oppositional defiant disorder | 236 | 30 |
| Panic | 5 | <1 |
| Psychosis spectrum symptoms | 226 | 29 |
| Posttraumatic stress disorder | 99 | 13 |
| Separation anxiety | 34 | 4 |
| Social phobia | 189 | 24 |
| Specific phobia | 242 | 31 |

<sup>a</sup> Because of comorbidity, many participants met the criteria for more than one disorder and were counted in multiple categories.
SD: standard deviation.

**Table S2.** A total of 108 psychopathology items contributed to the first latent component from partial least square correlation analysis.

| Num | Item Name | Weight (z) | Num | Item Name | Weight (z) | Num | Item Name | Weight (z) |
| --- | --- | --- | --- | --- | --- | --- | --- | --- |
| 1 | sip007 | 15.27 | 39 | add020 | 8.32 | 77 | ocd018 | 5.66 |
| 2 | sip011 | 12.50 | 40 | sip014 | 8.29 | 78 | odd005 | 5.62 |
| 3 | dep004 | 12.30 | 41 | cdd008 | 8.24 | 79 | sip028 | 5.32 |
| 4 | man007 | 12.10 | 42 | man005 | 7.93 | 80 | pan001 | 5.25 |
| 5 | sip003 | 11.91 | 43 | soc004 | 7.86 | 81 | ocd011 | 5.19 |
| 6 | sip013 | 11.67 | 44 | add014 | 7.80 | 82 | sip032 | 5.16 |
| 7 | sip005 | 11.43 | 45 | dep002 | 7.78 | 83 | agr002 | 5.02 |
| 8 | ocd005 | 11.30 | 46 | ocd013 | 7.73 | 84 | ptd009 | 4.99 |
| 9 | odd006 | 11.19 | 47 | sip004 | 7.72 | 85 | sep509 | 4.84 |
| 10 | soc005 | 10.69 | 48 | sui001 | 7.52 | 86 | sip038 | 4.80 |
| 11 | sip006 | 10.66 | 49 | soc002 | 7.52 | 87 | psy050 | 4.76 |
| 12 | agr005 | 10.60 | 50 | pan003 | 7.40 | 88 | sip039 | 4.54 |
| 13 | ocd004 | 10.40 | 51 | ocd002 | 7.33 | 89 | cdd005 | 4.52 |
| 14 | sip012 | 10.29 | 52 | phb002 | 7.29 | 90 | phb008 | 4.50 |
| 15 | odd002 | 10.14 | 53 | odd003 | 7.24 | 91 | cdd007 | 4.49 |
| 16 | sip009 | 9.94 | 54 | agr004 | 7.16 | 92 | ocd012 | 4.44 |
| 17 | ocd006 | 9.88 | 55 | phb006 | 7.11 | 93 | cdd002 | 4.41 |
| 18 | man002 | 9.85 | 56 | psy001 | 7.09 | 94 | ocd003 | 4.40 |
| 19 | agr001 | 9.85 | 57 | dep001 | 6.87 | 95 | cdd001 | 4.27 |
| 20 | dep006 | 9.59 | 58 | psy071 | 6.87 | 96 | sui002 | 4.22 |
| 21 | agr008 | 9.57 | 59 | sip033 | 6.77 | 97 | scr008 | 4.11 |
| 22 | agr006 | 9.52 | 60 | ocd008 | 6.75 | 98 | agr007 | 4.07 |
| 23 | man006 | 9.38 | 61 | ocd016 | 6.62 | 99 | agr003 | 4.04 |
| 24 | soc001 | 9.26 | 62 | ocd014 | 6.58 | 100 | gad001 | 3.83 |
| 25 | psy060 | 9.26 | 63 | add012 | 6.47 | 101 | sip027 | 3.81 |
| 26 | man004 | 9.23 | 64 | add021 | 6.44 | 102 | gad002 | 3.76 |
| 27 | sip010 | 9.10 | 65 | ocd001 | 6.25 | 103 | ocd007 | 3.62 |
| 28 | add016 | 9.06 | 66 | add022 | 6.15 | 104 | phb005 | 3.41 |
| 29 | phb004 | 8.99 | 67 | sep500 | 6.10 | 105 | scr001 | 3.35 |
| 30 | man001 | 8.95 | 68 | phb001 | 6.00 | 106 | cdd003 | 3.22 |
| 31 | phb007 | 8.73 | 69 | pan004 | 6.00 | 107 | ocd015 | 3.00 |
| 32 | add015 | 8.68 | 70 | sep510 | 5.87 | 108 | cdd006 | 2.86 |
| 33 | psy029 | 8.63 | 71 | sep508 | 5.84 | 109 | cdd004 | 1.76 |
| 34 | add011 | 8.61 | 72 | phb003 | 5.82 | 110 | scr007 | 1.33 |
| 35 | odd001 | 8.46 | 73 | psy070 | 5.81 | 111 | sep511 | 1.31 |
| 36 | man003 | 8.40 | 74 | add013 | 5.80 | 112 | scr006 | 1.11 |
| 37 | ocd019 | 8.36 | 75 | ocd017 | 5.76 |  |  |  |
| 38 | sip008 | 8.35 | 76 | soc003 | 5.69 |  |  |  |

The stability of contribution weight was evaluated by calculating the z-score (i.e., mean/SD) across repetitions. The four items that did not significantly contribute to the first component are marked with red color. See **Supplementary Data** for the question for each item.

## REFERENCES

Abdi, H. (2003). Partial least square regression (PLS regression). J Encyclopedia for research methods for the social sciences 6, 792–795.

Alexander-Bloch, A.F., Shou, H., Liu, S., Satterthwaite, T.D., Glahn, D.C., Shinohara, R.T., Vandekar, S.N., and Raznahan, A. (2018). On testing for spatial correspondence between maps of human brain structure and function. Neuroimage 178, 540–551.

Anderson, K.M., Ge, T., Kong, R., Patrick, L.M., Spreng, R.N., Sabuncu, M.R., Yeo, B.T.T., and Holmes, A.J. (2021). Heritability of individualized cortical network topography. Proc Natl Acad Sci U S A 118.

Avants, B.B., Tustison, N., and Song, G. (2009). Advanced normalization tools (ANTS). Insight j 2, 1–35.

Baker, J.T., Dillon, D.G., Patrick, L.M., Roffman, J.L., Brady, R.O., Jr., Pizzagalli, D.A., Ongur, D., and Holmes, A.J. (2019). Functional connectomics of affective and psychotic pathology. Proc Natl Acad Sci U S A 116, 9050–9059.

Barron, D.S., Gao, S., Dadashkarimi, J., Greene, A.S., Spann, M.N., Noble, S., Lake, E.M.R., Krystal, J.H., Constable, R.T., and Scheinost, D. (2020). Transdiagnostic, Connectome-Based Prediction of Memory Constructs Across Psychiatric Disorders. Cereb Cortex.

Bijsterbosch, J.D., Woolrich, M.W., Glasser, M.F., Robinson, E.C., Beckmann, C.F., Van Essen, D.C., Harrison, S.J., and Smith, S.M. (2018). The relationship between spatial configuration and functional connectivity of brain regions. Elife 7.

Braga, R.M., and Buckner, R.L. (2017). Parallel Interdigitated Distributed Networks within the Individual Estimated by Intrinsic Functional Connectivity. Neuron 95, 457–471 e455.

Cadwell, C.R., Bhaduri, A., Mostajo-Radji, M.A., Keefe, M.G., and Nowakowski, T.J. (2019). Development and Arealization of the Cerebral Cortex. Neuron 103, 980–1004.

Cai, D., He, X., Han, J., and Huang, T.S. (2011). Graph Regularized Nonnegative Matrix Factorization for Data Representation. IEEE Trans Pattern Anal Mach Intell 33, 1548–1560.

Calkins, M.E., Merikangas, K.R., Moore, T.M., Burstein, M., Behr, M.A., Satterthwaite, T.D., Ruparel, K., Wolf, D.H., Roalf, D.R., Mentch, F.D., et al. (2015). The Philadelphia Neurodevelopmental Cohort: constructing a deep phenotyping collaborative. Journal of child psychology and psychiatry, and allied disciplines 56, 1356–1369.

Calkins, M.E., Moore, T.M., Satterthwaite, T.D., Wolf, D.H., Turetsky, B.I., Roalf, D.R., Merikangas, K.R., Ruparel, K., Kohler, C.G., Gur, R.C., and Gur, R.E. (2017). Persistence of psychosis spectrum symptoms in the Philadelphia Neurodevelopmental Cohort: a prospective two-year follow-up. World Psychiatry 16, 62–76.

Casey, B.J., Cannonier, T., Conley, M.I., Cohen, A.O., Barch, D.M., Heitzeg, M.M., Soules, M.E., Teslovich, T., Dellarco, D.V., Garavan, H., et al. (2018). The Adolescent Brain Cognitive Development (ABCD) study: Imaging acquisition across 21 sites. Dev Cogn Neurosci 32, 43–54.

Caspi, A., Houts, R.M., Belsky, D.W., Goldman-Mellor, S.J., Harrington, H., Israel, S., Meier, M.H., Ramrakha, S., Shalev, I., Poulton, R., and Moffitt, T.E. (2014). The p Factor: One General Psychopathology Factor in the Structure of Psychiatric Disorders? Clin Psychol Sci 2, 119–137.

Caspi, A., and Moffitt, T.E. (2018). All for One and One for All: Mental Disorders in One Dimension. The American journal of psychiatry 175, 831–844.

Cholfin, J.A., and Rubenstein, J.L. (2007). Patterning of frontal cortex subdivisions by Fgf17. Proc Natl Acad Sci U S A 104, 7652–7657.

Ciric, R., Rosen, A.F.G., Erus, G., Cieslak, M., Adebimpe, A., Cook, P.A., Bassett, D.S., Davatzikos, C., Wolf, D.H., and Satterthwaite, T.D. (2018). Mitigating head motion artifact in functional connectivity MRI. Nat Protoc 13, 2801–2826.

Clark, D.A., Hicks, B.M., Angstadt, M., Rutherford, S., Taxali, A., Hyde, L., Weigard, A.S., Heitzeg, M.M., and Sripada, C. (2021). The general factor of psychopathology in the Adolescent Brain Cognitive Development (ABCD) Study: A comparison of alternative modeling approaches. Clinical Psychological Science 9, 169–182.

Clark, L.A., Cuthbert, B., Lewis-Fernandez, R., Narrow, W.E., and Reed, G.M. (2017). Three Approaches to Understanding and Classifying Mental Disorder: ICD-11, DSM-5, and the National Institute of Mental Health’s Research Domain Criteria (RDoC). Psychological science in the public interest : a journal of the American Psychological Society 18, 72–145.

Cole, M.W., Repovs, G., and Anticevic, A. (2014). The frontoparietal control system: a central role in mental health. Neuroscientist 20, 652–664.

Conway, C.C., Forbes, M.K., Forbush, K.T., Fried, E.I., Hallquist, M.N., Kotov, R., Mullins-Sweatt, S.N., Shackman, A.J., Skodol, A.E., South, S.C.*, et al.* (2019). A Hierarchical Taxonomy of Psychopathology Can Transform Mental Health Research. Perspect Psychol Sci 14, 419–436.

Cox, R.W. (1996). AFNI: software for analysis and visualization of functional magnetic resonance neuroimages. Computers and biomedical research, an international journal 29, 162–173.

Cui, Z., and Gong, G. (2018). The effect of machine learning regression algorithms and sample size on individualized behavioral prediction with functional connectivity features. Neuroimage 178, 622–637.

Cui, Z., Li, H., Xia, C.H., Larsen, B., Adebimpe, A., Baum, G.L., Cieslak, M., Gur, R.E., Gur, R.C., Moore, T.M., et al. (2020). Individual Variation in Functional Topography of Association Networks in Youth. Neuron 106, 340–353.

Drysdale, A.T., Grosenick, L., Downar, J., Dunlop, K., Mansouri, F., Meng, Y., Fetcho, R.N., Zebley, B., Oathes, D.J., Etkin, A., et al. (2017). Resting-state connectivity biomarkers define neurophysiological subtypes of depression. Nature medicine 23, 28–38.

Edition, F. (2013). Diagnostic and statistical manual of mental disorders. Am Psychiatric Assoc 21.

Elliott, M.L., Romer, A., Knodt, A.R., and Hariri, A.R. (2018). A Connectome-wide Functional Signature of Transdiagnostic Risk for Mental Illness. Biol Psychiatry 84, 452–459.

Fair, D.A., Schlaggar, B.L., Cohen, A.L., Miezin, F.M., Dosenbach, N.U., Wenger, K.K., Fox, M.D., Snyder, A.Z., Raichle, M.E., and Petersen, S.E. (2007). A method for using blocked and event-related fMRI data to study "resting state" functional connectivity. Neuroimage 35, 396–405.

Fischl, B. (2012). FreeSurfer. Neuroimage 62, 774–781.

Glasser, M.F., Coalson, T.S., Robinson, E.C., Hacker, C.D., Harwell, J., Yacoub, E., Ugurbil, K., Andersson, J., Beckmann, C.F., Jenkinson, M., et al. (2016). A multi-modal parcellation of human cerebral cortex. Nature 536, 171–178.

Gordon, E.M., Laumann, T.O., Adeyemo, B., Gilmore, A.W., Nelson, S.M., Dosenbach, N.U.F., and Petersen, S.E. (2017a). Individual-specific features of brain systems identified with resting state functional correlations. Neuroimage 146, 918–939.

Gordon, E.M., Laumann, T.O., Adeyemo, B., Huckins, J.F., Kelley, W.M., and Petersen, S.E. (2016). Generation and Evaluation of a Cortical Area Parcellation from Resting-State Correlations. Cereb Cortex 26, 288–303.

Gordon, E.M., Laumann, T.O., Adeyemo, B., and Petersen, S.E. (2017b). Individual Variability of the System-Level Organization of the Human Brain. Cereb Cortex 27, 386–399.

Gordon, E.M., Laumann, T.O., Gilmore, A.W., Newbold, D.J., Greene, D.J., Berg, J.J., Ortega, M., Hoyt-Drazen, C., Gratton, C., Sun, H., et al. (2017c). Precision Functional Mapping of Individual Human Brains. Neuron 95, 791–807 e797.

Gratton, C., Kraus, B.T., Greene, D.J., Gordon, E.M., Laumann, T.O., Nelson, S.M., Dosenbach, N.U., and Petersen, S.E. (2019). Defining Individual-Specific Functional Neuroanatomy for Precision Psychiatry. Biological Psychiatry.

Gratton, C., Laumann, T.O., Nielsen, A.N., Greene, D.J., Gordon, E.M., Gilmore, A.W., Nelson, S.M., Coalson, R.S., Snyder, A.Z., Schlaggar, B.L.*, et al.* (2018). Functional Brain Networks Are Dominated by Stable Group and Individual Factors, Not Cognitive or Daily Variation. Neuron 98, 439–452 e435.

Greene, D.J., Marek, S., Gordon, E.M., Siegel, J.S., Gratton, C., Laumann, T.O., Gilmore, A.W., Berg, J.J., Nguyen, A.L., Dierker, D.*, et al.* (2020). Integrative and Network-Specific Connectivity of the Basal Ganglia and Thalamus Defined in Individuals. Neuron 105, 742–758 e746.

Griffis, J.C., Metcalf, N.V., Corbetta, M., and Shulman, G.L. (2019). Structural disconnections explain brain network dysfunction after stroke. Cell reports 28, 2527–2540. e2529.

Hallquist, M.N., Hwang, K., and Luna, B. (2013). The nuisance of nuisance regression: spectral misspecification in a common approach to resting-state fMRI preprocessing reintroduces noise and obscures functional connectivity. Neuroimage 82, 208–225.

Haynes, J.D. (2015). A Primer on Pattern-Based Approaches to fMRI: Principles, Pitfalls, and Perspectives. Neuron 87, 257–270.

Insel, T., Cuthbert, B., Garvey, M., Heinssen, R., Pine, D.S., Quinn, K., Sanislow, C., and Wang, P. (2010). Research domain criteria (RDoC): toward a new classification framework for research on mental disorders. The American journal of psychiatry 167, 748–751.

Insel, T.R. (2014). The NIMH Research Domain Criteria (RDoC) Project: precision medicine for psychiatry. The American journal of psychiatry 171, 395–397.

Jenkinson, M., Beckmann, C.F., Behrens, T.E., Woolrich, M.W., and Smith, S.M. (2012). Fsl. Neuroimage 62, 782–790.

Kaczkurkin, A.N., Moore, T.M., Calkins, M.E., Ciric, R., Detre, J.A., Elliott, M.A., Foa, E.B., Garcia de la Garza, A., Roalf, D.R., Rosen, A., et al. (2018). Common and dissociable regional cerebral blood flow differences associate with dimensions of psychopathology across categorical diagnoses. Mol Psychiatry 23, 1981–1989.

Kaczkurkin, A.N., Moore, T.M., Sotiras, A., Xia, C.H., Shinohara, R.T., and Satterthwaite, T.D. (2020). Approaches to Defining Common and Dissociable Neurobiological Deficits Associated With Psychopathology in Youth. Biol Psychiatry 88, 51–62.

Kaczkurkin, A.N., Park, S.S., Sotiras, A., Moore, T.M., Calkins, M.E., Cieslak, M., Rosen, A.F.G., Ciric, R., Xia, C.H., Cui, Z., et al. (2019). Evidence for Dissociable Linkage of Dimensions of Psychopathology to Brain Structure in Youths. The American journal of psychiatry 176, 1000-1009.

Karcher, N.R., Michelini, G., Kotov, R., and Barch, D.M. (2020). Associations Between Resting-State Functional Connectivity and a Hierarchical Dimensional Structure of Psychopathology in Middle Childhood. Biol Psychiatry Cogn Neurosci Neuroimaging.

Karlaftis, V.M., Giorgio, J., Vertes, P.E., Wang, R., Shen, Y., Tino, P., Welchman, A.E., and Kourtzi, Z. (2019). Multimodal imaging of brain connectivity reveals predictors of individual decision strategy in statistical learning. Nat Hum Behav 3, 297–307.

Kaufman, J., Birmaher, B., Brent, D., Rao, U., Flynn, C., Moreci, P., Williamson, D., and Ryan, N. (1997). Schedule for Affective Disorders and Schizophrenia for School-Age Children-Present and Lifetime Version (K-SADS-PL): initial reliability and validity data. Journal of the American Academy of Child and Adolescent Psychiatry 36, 980–988.

Kaufmann, T., Alnaes, D., Doan, N.T., Brandt, C.L., Andreassen, O.A., and Westlye, L.T. (2017). Delayed stabilization and individualization in connectome development are related to psychiatric disorders. Nat Neurosci 20, 513–515.

Kebets, V., Holmes, A.J., Orban, C., Tang, S., Li, J., Sun, N., Kong, R., Poldrack, R.A., and Yeo, B.T.T. (2019). Somatosensory-Motor Dysconnectivity Spans Multiple Transdiagnostic Dimensions of Psychopathology. Biol Psychiatry 86, 779–791.

Kong, R., Li, J., Orban, C., Sabuncu, M.R., Liu, H., Schaefer, A., Sun, N., Zuo, X.N., Holmes, A.J., Eickhoff, S.B., and Yeo, B.T.T. (2019). Spatial Topography of Individual-Specific Cortical Networks Predicts Human Cognition, Personality, and Emotion. Cereb Cortex 29, 2533–2551.

Kotov, R., Krueger, R.F., Watson, D., Achenbach, T.M., Althoff, R.R., Bagby, R.M., Brown, T.A., Carpenter, W.T., Caspi, A., Clark, L.A., et al. (2017). The Hierarchical Taxonomy of Psychopathology (HiTOP): A dimensional alternative to traditional nosologies. Journal of abnormal psychology 126, 454–477.

Kotov, R., Krueger, R.F., Watson, D., Cicero, D.C., Conway, C.C., DeYoung, C.G., Eaton, N.R., Forbes, M.K., Hallquist, M.N., and Latzman, R.D. (2021). The Hierarchical Taxonomy of Psychopathology (HiTOP): A Quantitative Nosology Based on Consensus of Evidence. Annual Review of Clinical Psychology 17.

Krishnan, A., Williams, L.J., McIntosh, A.R., and Abdi, H. (2011). Partial Least Squares (PLS) methods for neuroimaging: a tutorial and review. Neuroimage 56, 455–475.

Krueger, R.F., Kotov, R., Watson, D., Forbes, M.K., Eaton, N.R., Ruggero, C.J., Simms, L.J., Widiger, T.A., Achenbach, T.M., Bach, B., et al. (2018). Progress in achieving quantitative classification of psychopathology. World Psychiatry 17, 282–293.

Lahey, B.B., Applegate, B., Hakes, J.K., Zald, D.H., Hariri, A.R., and Rathouz, P.J. (2012). Is there a general factor of prevalent psychopathology during adulthood? Journal of abnormal psychology 121, 971–977.

Lahey, B.B., Krueger, R.F., Rathouz, P.J., Waldman, I.D., and Zald, D.H. (2017). A hierarchical causal taxonomy of psychopathology across the life span. Psychol Bull 143, 142–186.

Lahey, B.B., Moore, T.M., Kaczkurkin, A.N., and Zald, D.H. (2021). Hierarchical models of psychopathology: empirical support, implications, and remaining issues. World Psychiatry 20, 57–63.

Larsen, B., and Luna, B. (2018). Adolescence as a neurobiological critical period for the development of higher-order cognition. Neuroscience and biobehavioral reviews 94, 179–195.

Laumann, T.O., Gordon, E.M., Adeyemo, B., Snyder, A.Z., Joo, S.J., Chen, M.Y., Gilmore, A.W., McDermott, K.B., Nelson, S.M., Dosenbach, N.U., et al. (2015). Functional System and Areal Organization of a Highly Sampled Individual Human Brain. Neuron 87, 657–670.

Lee, D.D., and Seung, H.S. (1999). Learning the parts of objects by non-negative matrix factorization. Nature 401, 788–791.

Li, H., Satterthwaite, T.D., and Fan, Y. (2017). Large-scale sparse functional networks from resting state fMRI. Neuroimage 156, 1–13.

Li, M., Wang, D., Ren, J., Langs, G., Stoecklein, S., Brennan, B.P., Lu, J., Chen, H., and Liu, H. (2019). Performing group-level functional image analyses based on homologous functional regions mapped in individuals. PLoS Biol 17, e2007032.

Lynch, C.J., Gunning, F.M., and Liston, C. (2020). Causes and Consequences of Diagnostic Heterogeneity in Depression: Paths to Discovering Novel Biological Depression Subtypes. Biol Psychiatry 88, 83–94.

Ma, Q., Tang, Y., Wang, F., Liao, X., Jiang, X., Wei, S., Mechelli, A., He, Y., and Xia, M. (2020). Transdiagnostic Dysfunctions in Brain Modules Across Patients with Schizophrenia, Bipolar Disorder, and Major Depressive Disorder: A Connectome-Based Study. Schizophr Bull 46, 699–712.

Marcus, D.S., Harms, M.P., Snyder, A.Z., Jenkinson, M., Wilson, J.A., Glasser, M.F., Barch, D.M., Archie, K.A., Burgess, G.C., Ramaratnam, M., et al. (2013). Human Connectome Project informatics: quality control, database services, and data visualization. Neuroimage 80, 202–219.

Margulies, D.S., Ghosh, S.S., Goulas, A., Falkiewicz, M., Huntenburg, J.M., Langs, G., Bezgin, G., Eickhoff, S.B., Castellanos, F.X., Petrides, M., et al. (2016). Situating the default-mode network along a principal gradient of macroscale cortical organization. Proc Natl Acad Sci U S A 113, 12574–12579.

Menon, V. (2011). Large-scale brain networks and psychopathology: a unifying triple network model. Trends Cogn Sci 15, 483–506.

Moore, T.M., Calkins, M.E., Satterthwaite, T.D., Roalf, D.R., Rosen, A.F.G., Gur, R.C., and Gur, R.E. (2019). Development of a computerized adaptive screening tool for overall psychopathology ("p"). J Psychiatr Res 116, 26–33.

Mourao-Miranda, J., Bokde, A.L., Born, C., Hampel, H., and Stetter, M. (2005). Classifying brain states and determining the discriminating activation patterns: Support Vector Machine on functional MRI data. Neuroimage 28, 980–995.

Mueller, S., Wang, D., Fox, M.D., Yeo, B.T., Sepulcre, J., Sabuncu, M.R., Shafee, R., Lu, J., and Liu, H. (2013). Individual variability in functional connectivity architecture of the human brain. Neuron 77, 586–595.

Muthen, B., and Asparouhov, T. (2012). Bayesian structural equation modeling: a more flexible representation of substantive theory. Psychol Methods 17, 313–335.

Norman, K.A., Polyn, S.M., Detre, G.J., and Haxby, J.V. (2006). Beyond mind-reading: multi-voxel pattern analysis of fMRI data. Trends Cogn Sci 10, 424–430.

O’Leary, D.D., Chou, S.J., and Sahara, S. (2007). Area patterning of the mammalian cortex. Neuron 56, 252–269.

Ojemann, J.G., Akbudak, E., Snyder, A.Z., McKinstry, R.C., Raichle, M.E., and Conturo, T.E. (1997). Anatomic localization and quantitative analysis of gradient refocused echo-planar fMRI susceptibility artifacts. Neuroimage 6, 156–167.

Pedregosa, F., Varoquaux, G., Gramfort, A., Michel, V., Thirion, B., Grisel, O., Blondel, M., Prettenhofer, P., Weiss, R., and Dubourg, V. (2011). Scikit-learn: Machine learning in Python. the Journal of machine Learning research 12, 2825–2830.

Power, J.D., Cohen, A.L., Nelson, S.M., Wig, G.S., Barnes, K.A., Church, J.A., Vogel, A.C., Laumann, T.O., Miezin, F.M., Schlaggar, B.L., and Petersen, S.E. (2011). Functional network organization of the human brain. Neuron 72, 665–678.

Qi, S., Bustillo, J., Turner, J.A., Jiang, R., Zhi, D., Fu, Z., Deramus, T.P., Vergara, V., Ma, X., Yang, X.*, et al.* (2020). The relevance of transdiagnostic shared networks to the severity of symptoms and cognitive deficits in schizophrenia: a multimodal brain imaging fusion study. Transl Psychiatry 10, 149.

Romer, A.L., Elliott, M.L., Knodt, A.R., Sison, M.L., Ireland, D., Houts, R., Ramrakha, S., Poulton, R., Keenan, R., Melzer, T.R., et al. (2021). Pervasively Thinner Neocortex as a Transdiagnostic Feature of General Psychopathology. The American journal of psychiatry 178, 174–182.

Satterthwaite, T.D., Elliott, M.A., Gerraty, R.T., Ruparel, K., Loughead, J., Calkins, M.E., Eickhoff, S.B., Hakonarson, H., Gur, R.C., Gur, R.E., and Wolf, D.H. (2013a). An improved framework for confound regression and filtering for control of motion artifact in the preprocessing of resting-state functional connectivity data. Neuroimage 64, 240–256.

Satterthwaite, T.D., Elliott, M.A., Ruparel, K., Loughead, J., Prabhakaran, K., Calkins, M.E., Hopson, R., Jackson, C., Keefe, J., Riley, M., et al. (2014). Neuroimaging of the Philadelphia neurodevelopmental cohort. Neuroimage 86, 544–553.

Satterthwaite, T.D., Feczko, E., Kaczkurkin, A.N., and Fair, D.A. (2020). Parsing Psychiatric Heterogeneity Through Common and Unique Circuit-Level Deficits. Biol Psychiatry 88, 4–5.

Satterthwaite, T.D., Wolf, D.H., Erus, G., Ruparel, K., Elliott, M.A., Gennatas, E.D., Hopson, R., Jackson, C., Prabhakaran, K., Bilker, W.B., et al. (2013b). Functional maturation of the executive system during adolescence. J Neurosci 33, 16249–16261.

Sha, Z., Wager, T.D., Mechelli, A., and He, Y. (2019). Common Dysfunction of Large-Scale Neurocognitive Networks Across Psychiatric Disorders. Biol Psychiatry 85, 379–388.

Shanmugan, S., Wolf, D.H., Calkins, M.E., Moore, T.M., Ruparel, K., Hopson, R.D., Vandekar, S.N., Roalf, D.R., Elliott, M.A., Jackson, C., et al. (2016). Common and Dissociable Mechanisms of Executive System Dysfunction Across Psychiatric Disorders in Youth. Am J Psychiatry 173, 517–526.

Sheffield, J.M., Kandala, S., Tamminga, C.A., Pearlson, G.D., Keshavan, M.S., Sweeney, J.A., Clementz, B.A., Lerman-Sinkoff, D.B., Hill, S.K., and Barch, D.M. (2017). Transdiagnostic Associations Between Functional Brain Network Integrity and Cognition. JAMA Psychiatry 74, 605–613.

Smith, S.M., Jenkinson, M., Woolrich, M.W., Beckmann, C.F., Behrens, T.E., Johansen-Berg, H., Bannister, P.R., De Luca, M., Drobnjak, I., Flitney, D.E., et al. (2004). Advances in functional and structural MR image analysis and implementation as FSL. Neuroimage 23 Suppl 1, S208–219.

Somerville, L.H., Bookheimer, S.Y., Buckner, R.L., Burgess, G.C., Curtiss, S.W., Dapretto, M., Elam, J.S., Gaffrey, M.S., Harms, M.P., Hodge, C., et al. (2018). The Lifespan Human Connectome Project in Development: A large-scale study of brain connectivity development in 5-21 year olds. Neuroimage 183, 456–468.

Sotiras, A., Toledo, J.B., Gur, R.E., Gur, R.C., Satterthwaite, T.D., and Davatzikos, C. (2017). Patterns of coordinated cortical remodeling during adolescence and their associations with functional specialization and evolutionary expansion. Proc Natl Acad Sci U S A 114, 3527–3532.

Sydnor, V.J., Larsen, B., Bassett, D., Alexander-Bloch, A., Fair, D.A., Liston, C., Mackey, A.P., Milham, M.P., Pines, A.R., Roalf, D.R., et al. (2021). Neurodevelopment of the association cortices: Patterns, mechanisms, and implications for psychopathology. Neuron.

Sylvester, C.M., Yu, Q., Srivastava, A.B., Marek, S., Zheng, A., Alexopoulos, D., Smyser, C.D., Shimony, J.S., Ortega, M., Dierker, D.L., et al. (2020). Individual-specific functional connectivity of the amygdala: A substrate for precision psychiatry. Proc Natl Acad Sci U S A 117, 3808–3818.

Vandekar, S.N., Shinohara, R.T., Raznahan, A., Roalf, D.R., Ross, M., DeLeo, N., Ruparel, K., Verma, R., Wolf, D.H., and Gur, R.C. (2015). Topologically dissociable patterns of development of the human cerebral cortex. Journal of Neuroscience 35, 599–609.

Wang, D., Buckner, R.L., Fox, M.D., Holt, D.J., Holmes, A.J., Stoecklein, S., Langs, G., Pan, R., Qian, T., Li, K.*, et al.* (2015). Parcellating cortical functional networks in individuals. Nat Neurosci 18, 1853–1860.

Wang, D., Li, M., Wang, M., Schoeppe, F., Ren, J., Chen, H., Ongur, D., Baker, J.T., and Liu, H. (2018). Individual-specific functional connectivity markers track dimensional and categorical features of psychotic illness. Mol Psychiatry.

Wig, G.S., Laumann, T.O., and Petersen, S.E. (2014). An approach for parcellating human cortical areas using resting-state correlations. Neuroimage 93 Pt 2, 276–291.

Wolf, D.H., Satterthwaite, T.D., Calkins, M.E., Ruparel, K., Elliott, M.A., Hopson, R.D., Jackson, C.T., Prabhakaran, K., Bilker, W.B., Hakonarson, H.*, et al.* (2015). Functional neuroimaging abnormalities in youth with psychosis spectrum symptoms. JAMA Psychiatry 72, 456–465.

Xia, C.H., Ma, Z., Ciric, R., Gu, S., Betzel, R.F., Kaczkurkin, A.N., Calkins, M.E., Cook, P.A., Garcia de la Garza, A., Vandekar, S.N., et al. (2018). Linked dimensions of psychopathology and connectivity in functional brain networks. Nat Commun 9, 3003.

Yeo, B.T., Krienen, F.M., Sepulcre, J., Sabuncu, M.R., Lashkari, D., Hollinshead, M., Roffman, J.L., Smoller, J.W., Zollei, L., Polimeni, J.R.*, et al.* (2011). The organization of the human cerebral cortex estimated by intrinsic functional connectivity. J Neurophysiol 106, 1125–1165.

Yoo, K., Rosenberg, M.D., Hsu, W.T., Zhang, S., Li, C.R., Scheinost, D., Constable, R.T., and Chun, M.M. (2018). Connectome-based predictive modeling of attention: Comparing different functional connectivity features and prediction methods across datasets. Neuroimage 167, 11–22.

